# Evolution of the substrate specificity of an RNA ligase ribozyme from phosphorimidazole- to triphosphate-activation

**DOI:** 10.1101/2024.04.11.588910

**Authors:** Saurja DasGupta, Zoe Weiss, Collin Nisler, Jack W. Szostak

## Abstract

The acquisition of new RNA functions through evolutionary processes would have been essential for the diversification of RNA-based primordial biology and its subsequent transition to modern biology. However, the mechanisms by which RNAs access new functions remain unclear. Do ribozymes need completely new folds to support new but related functions, or is re-optimization of the active site sufficient? What are the roles of neutral and adaptive mutations in evolutionary innovation? Here we address these questions experimentally by focusing on the evolution of substrate specificity in RNA-catalyzed RNA assembly reactions. We use directed *in vitro* evolution to show that a ligase ribozyme that uses prebiotically relevant 5′-phosphorimidazole-activated substrates can be evolved to catalyze ligation with substrates that are 5′-activated with the biologically relevant triphosphate group. Interestingly, despite catalyzing a related reaction, the new ribozyme folds into a completely new structure and exhibits promiscuity by catalyzing RNA ligation with both triphosphate and phosphorimidazole-activated substrates. Although distinct in sequence and structure, the parent phosphorimidazolide ligase and the evolved triphosphate ligase ribozymes can be connected by a series of point mutations where the intermediate sequences retain at least some ligase activity. The existence of a quasi-neutral pathway between these distinct ligase ribozymes suggests that neutral drift is sufficient to enable the acquisition of new substrate specificity, thereby providing opportunities for subsequent adaptive optimization. The transition from RNA-catalyzed RNA assembly using phosphorimidazole-activated substrates to triphosphate-activated substrates may have set the stage for the later evolution of the protein enzymes that use monomeric triphosphates (NTPs) for RNA synthesis.

## INTRODUCTION

In an early stage of the evolution of life, referred to as the RNA World, primordial cells are thought to have used RNA to constitute both genomes and enzymes (ribozymes). In the absence of material remains from the RNA World, experimental models are used to understand the various chemical and biochemical processes that would have been important for the origin and evolution of early life. As efficient RNA assembly is a prerequisite for the replication of genomic and catalytic RNAs, there has been considerable focus on modeling RNA ligation and polymerization pathways in the absence and presence of enzymes. Protein enzymes (RNA polymerases) use nucleoside triphosphates (NTPs) as substrates for the synthesis of RNA. However, the low reactivity of 5′-triphosphorylated monomers/oligomers for nonenzymatic polymerization/ligation suggests that before the emergence of enzymes, RNA building blocks were activated with more reactive groups. 5′-phosphorimidazole-activated monomers and oligomers have been widely used to study nonenzymatic RNA assembly because of their intrinsic reactivity.(1–4) Recent discoveries provide strong evidence for the prebiotic relevance of phosphorimidazolides.(5–8) To model the crucial transition from nonenzymatic RNA assembly to ribozyme-catalyzed RNA assembly, we previously reported the *in vitro* selection of ribozymes that use 5′-phosphorimidazolide RNA as substrates for ligation.(9)

Although beneficial for both enzyme-free and ribozyme-catalyzed RNA assembly, the enhanced reactivity of phosphorimidazolides makes them susceptible to hydrolysis. Therefore, kinetically more stable 5′-triphosphorylated building blocks would have been better substrates for RNA assembly if they were readily generated on the early Earth and suitable enzymes were available to use them. 5′-triphosphorylated RNA monomers and oligomers can be generated by reactions with abiotically synthesized polyphosphates, such as cyclic-trimetaphosphate(10–12) and ribozymes that accelerate this triphosphorylation reaction have been reported.(13, 14) Additionally, triphosphate substrates may be produced by the reaction of phosphorimidazolides with pyrophosphate, the latter abundantly generated in mineral films in alkaline hydrothermal environments.(15–17) As the uptake of trimetaphosphate or triphosphorylated RNA oligonucleotide substrates by primitive cells seems unlikely, given that even NTPs show low permeability to fatty acid vesicles,(18) triphosphorylated substrates could plausibly be generated by the intracellular conversion of phosphorimidazolides to triphosphates through reactions with encapsulated pyrophosphate.

The availability of 5′-triphosphorylated building blocks would impart a selective advantage to ribozymes that used such substrates. Multiple ribozymes that use such substrates have been artificially evolved.(3, 19–27) But how, in the RNA World, might ribozymes that used phosphorimidazolides have acquired the ability to use triphosphates? The chemistry of RNA assembly with these two substrates differs only in the identity of the leaving group (imidazole vs. pyrophosphate). Would a minor reorganization of the active site allow ligases to switch from phosphorimidazolides to triphosphates, or is a completely new catalytic fold required? Results from RNA evolution experiments where new functions were evolved from existing ones indicate that sequences often need to diverge greatly to access new functions.(28–33) This observation is intriguing as RNA fitness landscapes are considered to be rugged and composed of isolated high-activity peaks separated by low-activity valleys.(34–38) As evolutionary adaptation is conceptualized as the climbing of fitness peaks, adaptation across rugged landscapes is expected to be difficult. However, functional RNAs can be robust to mutational perturbation, resulting in the existence of mutational pathways composed of functional sequence variants (‘neutral pathways’) between fitness peaks.(39, 40) Therefore, new RNA functions may be accessed via neutral or quasi-neutral drifts across the fitness landscape followed by adaptive optimization (i.e., peak climbing). Although it has been suggested that neutral networks pervade the RNA sequence space and connect distinct phenotypes, (41–43) only a handful of examples have been reported.(37, 43, 44) Understanding the role of neutral pathways in the evolution of new functions in RNA is crucial for explaining the evolutionary diversification of ribozymes and, consequently, the emergence of RNA-based biology.

Here, we report the *in vitro* evolution of a new ligase that uses 5′-triphosphorylated substrates (‘PPP-ligase’), starting from a previously characterized ligase ribozyme that uses 5′-phosphoro-2-aminoimidazolide activated RNA oligonucleotides as substrates (‘AIP-ligase’). The PPP-ligase ribozyme differs from the parent AIP-ligase by 28 mutations and consequently adopts a new structure. The new PPP-ligase, in addition to catalyzing the ligation of triphosphorylated substrates, also catalyzes the ligation of phosphorimidazolide substrates, making it catalytically promiscuous. Despite being separated by 28 mutations, the parent AIP-ligase and the newly evolved PPP-ligase are connected by a quasi-neutral pathway where each point mutant catalyzes ligation with either phosphorimidazolide or both phosphorimidazolide and triphosphate substrates. Promiscuous ribozymes with the capacity to catalyze RNA assembly using both prebiotically-relevant phosphorimidazolides and biologically-relevant triphosphates could have led to the evolution of enzymatic pathways for the synthesis of triphosphate substrates. This would, in turn, have set the stage for the later emergence of protein polymerases that use the same triphosphate substrate activation that is now universal in biology.

## RESULTS

### Isolation of new ligase ribozymes through directed evolution

In order to select new ligase ribozymes that use 5′-triphosphorylated substrates, we prepared a sequence library derived from a previously characterized AIP-ligase ribozyme (hereafter, RS1) that comprised a 40 nt catalytic domain flanked by an 8 base-pair stem (Fig. 1, Fig. 2A). The catalytic domain was partially randomized at 21% per position (i.e., 7% each of the three non-RS1 nucleotides) to generate the selection library (wild type RS1 = 0.008%). The library was connected to an 8 nt primer sequence at its 3′ end via a U6 linker. The constant 5′ and 3′ regions contained primer binding sites for PCR amplification. We started with 3 nmol of the RNA library transcribed from 1.2 nmol dsDNA (estimated complexity of ∼10^15^ sequences). The library was challenged with a 5′-triphosphorylated RNA oligonucleotide substrate that was biotinylated at its 3′ end (‘PPP-LigB’). An external RNA template was supplied to bring the primer (i.e. 3′-end of the library RNAs) and the substrate into close proximity, to facilitate the targeted reaction between the primer 3′-OH and the substrate 5′-α-phosphate (Fig. 1). Active library sequences were affinity-purified using streptavidin-coated magnetic beads, by virtue of their covalent attachment to the biotin-tagged substrate. The captured sequences were selectively amplified by RT-PCR using appropriate primers (Table S6) and *in vitro* transcribed to generate an enriched library for the next round of selection (Fig. 1). Selection stringency was increased by reducing the reaction time and/or lowering the Mg^2+^ concentration in each round to isolate the most active ribozymes (Table S1).

We assayed ligase activities of the output library from each round and observed significant activity after round 4 (Fig. 2B). After round 6, the library was found to ligate the selection substrate, PPP-LigB, three orders of magnitude faster than the starting library. In contrast, both the round 6 library and the starting library exhibited comparable ligase activities toward an identical RNA sequence with a 5′-phosphoro-2-aminoimidazolide group (‘AIP-Lig’) (Fig. 2B). Surprisingly, the starting library, generated by partially mutagenizing an active AIP-ligase, (Fig. 2A) demonstrated low but detectable triphosphate ligation (*k*obs = 0.00094 ± 0.0001/h), while no ligation was detected after 3 days with a completely randomized library. This suggests that even the initial doped library is able to catalyze extremely slow ligation with the triphosphate substrate, PPP-LigB.

High-throughput sequencing of the output from each round of selection revealed significant enrichment of specific sequences after round 4, with the three most abundant sequences comprising ∼60% of the total reads in round 6 (Fig. 2C, Table 1). Early rounds of selection showed a small increase in the representation of the parental AIP-ligase, RS1 (Fig. 2C), from an expected abundance of ∼0.01% in the initial doped library to 0.025% after round 2 and round 3, after which the parental sequence exhibits a sharp decline in abundance. Further rounds of selection led to continuing decrease in the sequence diversity of the sequence library (Table S1). The decline in both the abundance of RS1 and the overall sequence diversity, together with the emergence of new dominant sequences, collectively suggested the isolation of new ligase ribozymes.

**Fig. 1.**
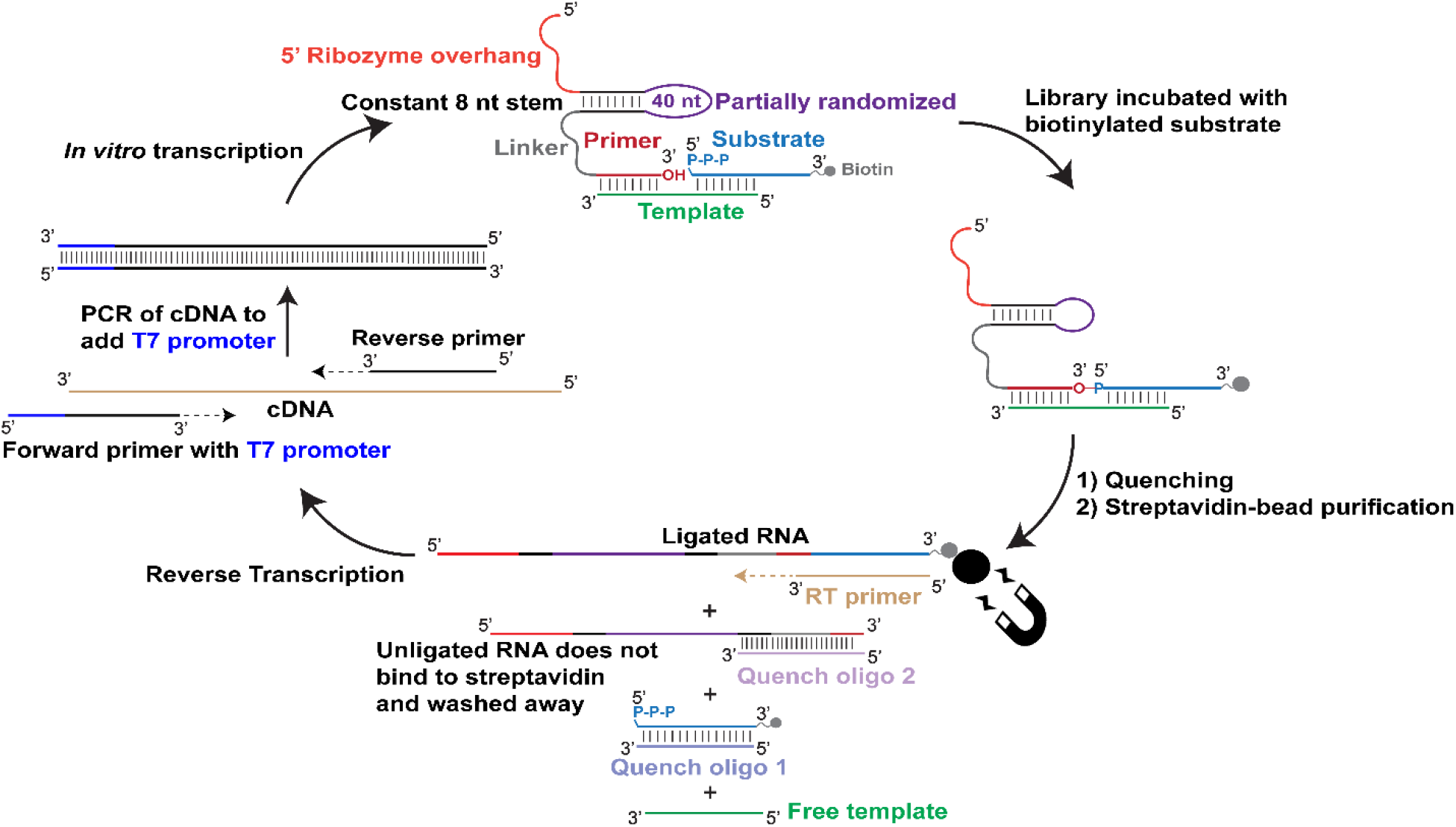
Selection strategy to isolate ribozymes that catalyze the ligation of 5′-triphosphorylated RNA. A partially randomized library derived from an existing AIP-ligase was incubated with an RNA template and a 5′-triphosphorylated RNA substrate that was biotinylated at its 3′ end. Reactions were quenched by adding EDTA. Excess DNA oligos complementary to the substrate and the 3′ ends of unligated library sequences were added to disrupt the ternary complex formed by the library, template, and substrate sequences, thus preventing the retention of inactive sequences on beads due to noncovalent association with the biotinylated substrate. Sequences that catalyzed ligation were captured on streptavidin-coated magnetic beads and reverse transcribed using a primer that is complementary to the substrate, thus forcing the selection of the desired ligases. The resulting cDNA was converted to dsDNA and amplified by PCR, which also added a T7 promoter sequence to the dsDNA. The dsDNA was *in vitro* transcribed to generate the RNA library for the next round of selection.

### A single ribozyme class catalyzes the ligation of 5′-triphosphorylated RNA

Sequences isolated from round 6 were sorted into 13 distinct clusters of closely-related sequences with little sequence overlap between clusters. Sequences in the three most abundant clusters comprised 92% of all sequence reads, and the dominant sequences (hereafter, peak sequences) of each cluster represented 50-80% of all reads within their clusters (Table 1). To capture the diversity of the selected ligases, all peak sequences that occupied >0.1% of the total sequence reads were tested for their ability to ligate to the selection substrate, PPP-LigB, in addition to a triphosphorylated substrate that lacked a biotin tag (PPP-Lig) and the phosphorimidazolide substrate, AIP-Lig. Peak sequences from CS1-CS5, representing the five most abundant ligase clusters, satisfied this threshold. While CS3 catalyzed ligation with all three substrates, CS1, CS2, CS4, and CS5 catalyzed ligation with only PPP-LigB (Fig. 2D). The failure to ligate PPP-Lig suggests that CS1, CS2, CS4, and CS5 do not catalyze the desired reaction, but survived selection by using alternative ligation mechanism(s) (a detailed analysis will be reported elsewhere). On the other hand, CS3’s ability to ligate both phosphorimidazolide and triphosphate substrates indicated that we had isolated a promiscuous ligase.

**Fig. 2.**
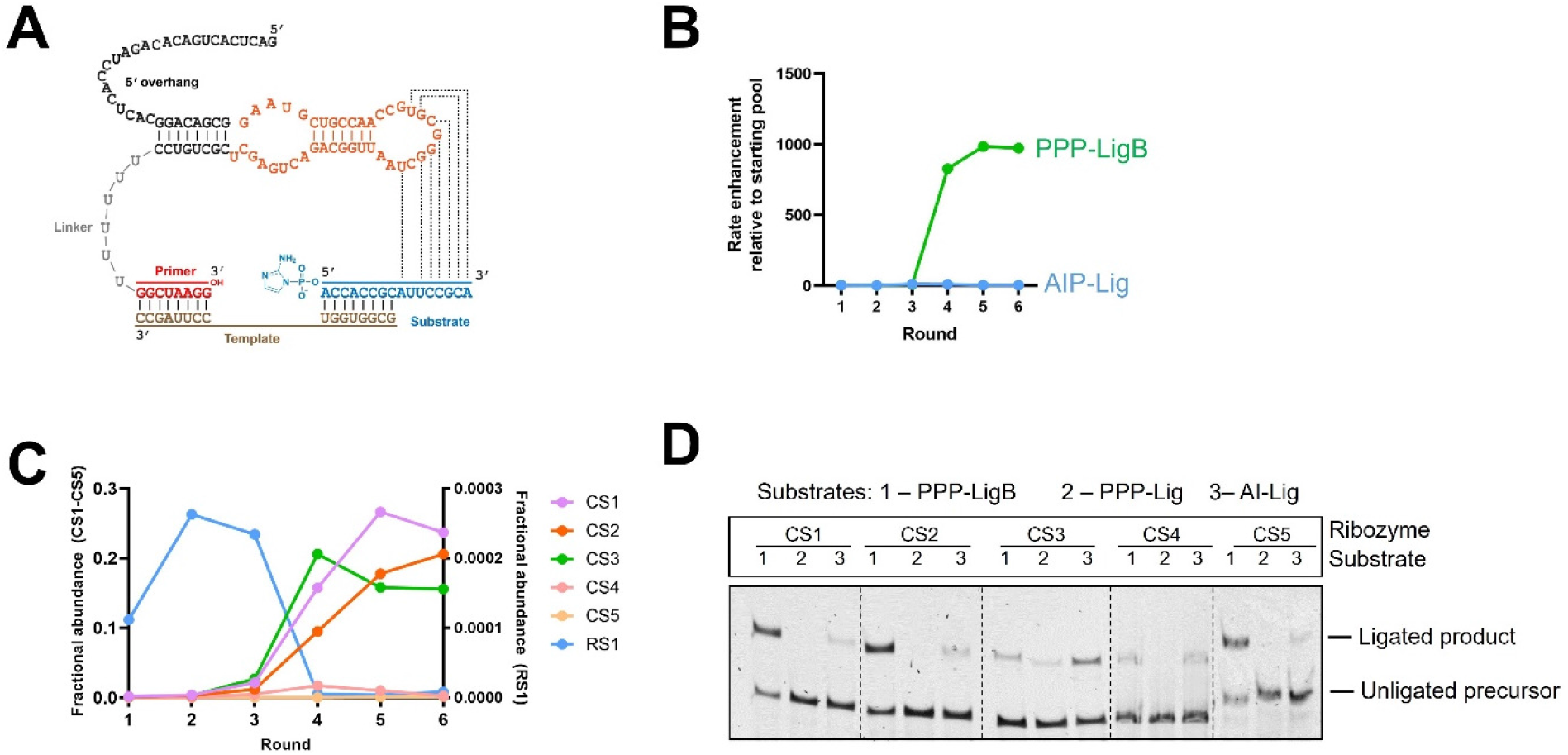
Enrichment of catalytically active sequences during selection and their biochemical validation. **A.** Secondary structure of the parent AIP-ligase ribozyme that was partially mutagenized to generate the selection library. The library region is highlighted in orange. Dashed lines indicate putative base-pairing interactions between the ribozyme and the substrate. **B.** Significant enrichment in activity (∼1000-fold) was observed after round 4, which was maintained through round 6. **C.** Certain sequences became predominant after round 4, which coincided with the abrupt fall in the relative abundance of the parent AIP-ligase, RS1. Fractional abundances of CS1-CS5 and RS1 are shown in the left and right y-axes, respectively. **D.** Ligase activities of the five most abundant sequences with (1) 5′ -riphosphorylated and 3′ biotinylated substrate (PPP-LigB), (2) 5′-triphosphorylated substrate (PPP-Lig), and (3) 5′-phosphorimidazolide substrate (AIP-Lig). Only CS3 is able to ligate PPP-Lig and AI-Lig. Ligation reactions contained 1 µM ribozyme, 1.2 µM RNA template, and 2 µM RNA substrate in 100 mM Tris-HCl pH 8.0, 300 mM NaCl, and either 10 mM MgCl2 (for ligation with AIP-Lig) or 100 mM MgCl2 (for ligation with PPP-Lig/PPP-LigB).

**Table 1.**
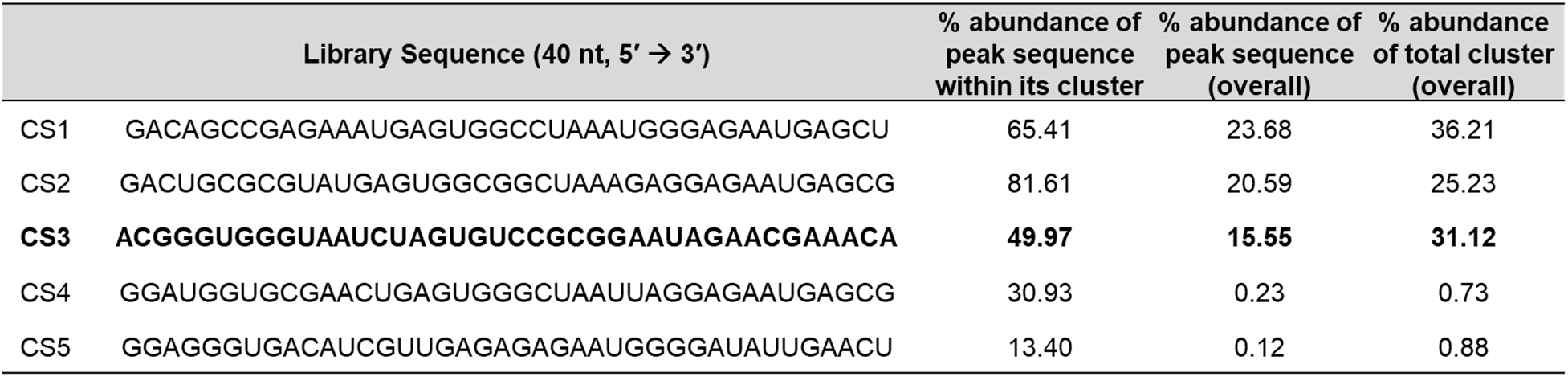
The five most abundant sequences identified by high-throughput sequencing. Sequences CS1-CS5 represent the most abundant sequences in their respective clusters (referred to as peak sequences). While the peak sequences in clusters CS1-CS5 occupied ∼60% of the round 6 population, the clusters as a whole accounted for >90% of all sequences. The PPP-ligase, CS3, which occupied ∼15% of the round 6 population, is shown in boldface. The 40 nt library sequences shown here constitute the partially randomized region highlighted in orange in Fig. 2A.

### Ligation requires a template and generates a 3′-5′ phosphodiester linkage

To confirm that CS3 catalyzes the desired reaction, we first tested substrates with different 5′ chemistries, with or without a template (Fig. 3A, B). Ligation was eliminated in the absence of the template. Substrates with unactivated 5′-monophosphates, with or without 3′ biotin (P-B or P) failed to undergo ligation, while 5′-triphosphate substrates were ligated by CS3. CS3 also ligated to a substrate with 5′-phosphorimidazole activation (AIP). Next, we tested CS3 truncation constructs in which either the first 25 nt, consisting of the PCR primer binding sequence (5′t) or the last 14 nt, including the 3′ primer (3′t), were deleted. Both truncations abolished ligation (Fig. 3C). As the 3′ truncation involves the deletion of a presumably unstructured region (U6 linker + 8 nt primer), loss of activity due to ribozyme misfolding is unlikely. This implicated the 3′ primer sequence in CS3-catalyzed ligation. The deleted 5′ sequence in 5′t, on the other hand, base-pairs extensively with the central 40 nt region corresponding to the mutagenized segment (Fig. 3A) (See Fig. S5 for secondary structure determination by SHAPE probing), explaining the loss of activity upon 5′ truncation. Previous selections targeting 3′-5′ ligation have resulted in the isolation of ribozymes that catalyze ligation between the ribozyme 5′-triphosphate and the substrate 5′-phosphorimidazole group.(45) To test this possibility, we examined the activities of CS3 variants containing either a 5′-monophosphate (5′P) or a 5′-hydroxyl (5′OH) moiety. As expected, both CS3 variants exhibited PPP-ligation, consistent with ligation at the 3′ end of the ribozyme (Fig. 3D).

**Fig. 3.**
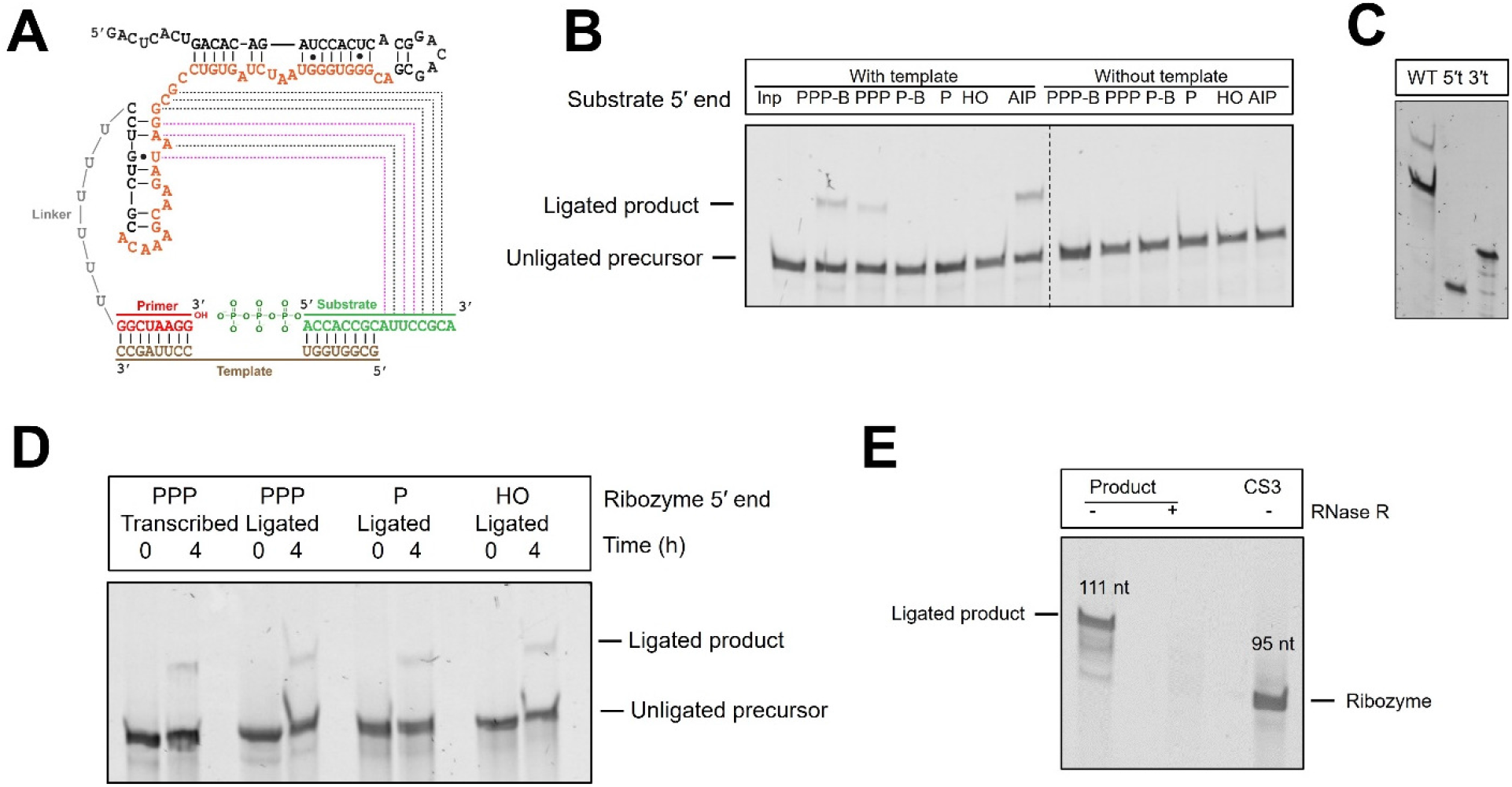
CS3-catalyzed templated ligation to triphosphate substrates involves the 3′-hydroxyl group of the primer and the 5′-triphosphate group of the substrate. **A.** SHAPE-derived secondary structure of CS3 in complex with the template and substrate RNA sequences (See Fig. S5 for SHAPE data). The region mutagenized in the library is highlighted in orange. Dashed lines indicate putative base-pairing interactions between the ribozyme and the substrate (see below). Black and magenta lines indicate complementarity between the substrate 3′ end and the unpaired and paired residues of the ribozyme, respectively. **B.** CS3-catalyzed RNA ligation requires a template and 5′ substrate activation, either in the form of triphosphate or phosphorimidazole. **C.** Truncation of CS3 by deleting the first 25 nt from the 5′ end or the last 14 nucleotides from the 3′ end results in the loss of activity. **D.** CS3 tolerates changes to its 5′ chemistry indicating that its 5′ end does not participate in ligation. **E.** The 111 nt product of ligation between CS3 and PPP-Lig is completely degraded by RNase R, which suggests that it contains a 3′-5′ linkage between the ribozyme and substrate. The product lanes contain purified ligated product, incubated with or without RNase R. Ligation reactions contained 1 µM ribozyme, 1.2 µM RNA template (in templated reactions), and 2 µM RNA substrate in 100 mM Tris-HCl pH 8.0, 300 mM NaCl, and either 10 mM MgCl2 (for ligation with AIP-Lig) or 100 mM MgCl2 (for ligation with PPP-Lig/PPP-LigB).

With strong support for the participation of the ribozyme 3′ end and substrate 5′ end, we investigated the importance of base-pairing at the ligation junction by mutating the relevant template nucleotides. A C to G mutation that disrupts a base-pair between the 3′ nucleotide of the primer and the template preserved ligation, but a U to A mutation that disrupts an A-U base-pair between the 5′ end of the substrate and the template, abrogated ligation (Fig. S1). The parent AIP-ligase, RS1, generates a 3′-5′ linkage, and 3′-5′ linkages form the canonical backbone of modern RNA. To characterize the linkage created as a result of CS3-catalyzed ligation with PPP-Lig, we purified the ligated product and incubated it with RNase R, a 3′◊5′ exonuclease. We observed complete degradation of the ligated product suggesting the presence of a natural 3′-′5′ linkage as opposed to a 2′-5′ linkage, which would result in exonuclease stalling at the linkage (Fig. 3E). Collectively, these results confirmed that CS3-catalyzes the desired reaction, i.e., templated ligation of 5′-triphosphorylated RNA generating a 3′-5′ phosphodiester linkage between the ribozyme and the substrate.

### Comparison with other ribozyme-catalyzed ligation reactions

Template-directed nonenzymatic RNA ligation has been explored for the last 50 years as a model for prebiotic RNA assembly.(46, 47) Imidazoles are superior leaving groups to pyrophosphates, therefore, templated phosphorimidazolide ligation, unlike ligation with triphosphorylated RNA, is detectable even in the absence of enzymes. For example, at pH 8 and 100 mM Mg^2+^, *k*obs for template-directed nonenzymatic ligation with triphosphorylated RNA is ∼1.3 x 10^-5^ h^-1^, while the corresponding *k*obs with a 2AI-activated RNA is 2.4 x 10^-2^ h^-1^ approximately three orders of magnitude faster.(9, 47) While previously isolated AIP-ligases exhibit rate enhancements of 170-740-fold (at 10 mM Mg^2+^),(9) the fastest PPP-ligases accelerate ligation by 10^6^-fold (at 100 mM Mg^2+^).(19, 21, 48, 49) CS3, although specifically selected for PPP-ligation, also catalyzes AIP-ligation (Fig. 4A). CS3 accelerates PPP-ligation by ∼10^3^-fold at 100 mM Mg^2+^, and AIP-ligation by ∼100-fold at 10 mM Mg^2+^, (Fig. 4B), making it slower than most of the previously reported PPP-ligases and AIP-ligases (Fig. 4C).

**Fig. 4.**
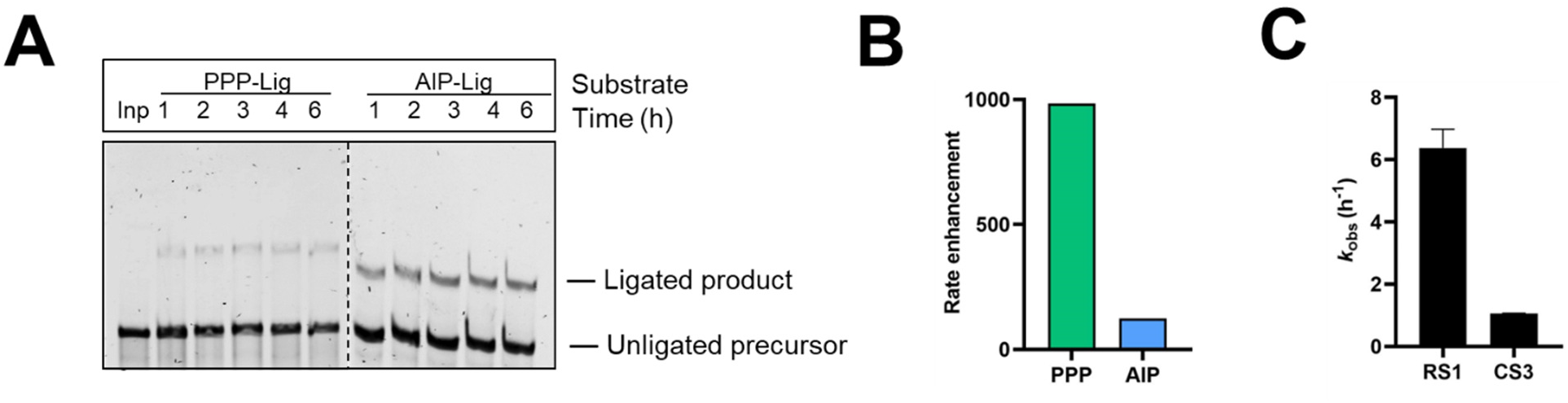
CS3 catalyzes the ligation of both triphosphate and phosphorimidazolide RNA substrates. **A.** Representative time course for CS3-catalyzed ligation of PPP-and AIP-substrates. **B.** CS3 accelerates PPP-ligation by ∼1000-fold and AIP-ligation by ∼100-fold relative to their corresponding background reactions. **C.** CS3 is ∼6-fold slower than the parent AIP-ligase, RS1. Ligation reactions contained 1 µM ribozyme, 1.2 µM RNA template, and 2 µM RNA substrate (AIP-Lig or PPP-Lig) in 100 mM Tris-HCl pH 8.0, 300 mM NaCl, and either 10 mM MgCl2 (AIP-Lig) or 100 mM MgCl2 (PPP-Lig).

Ribozymes often increase reaction rates by tightly binding and positioning catalytic Mg^2+^ ion(s) within the active site.(9, 19, 48, 50) The first ligase ribozyme reported to use triphosphorylated RNA substrates (‘class I’ ligase) uses Mg^2+^ for neutralizing the negative charge on the pyrophosphate group and activating the 3′-O nucleophile,(51) strategies also used by protein-based polymerases.(52–56) A Mg^2+^-titration curve for CS3-catalyzed PPP-ligation revealed a sharp increase in *k*obs between 5 mM and 25 mM followed by a plateau of the rate constant with increasing Mg^2+^ concentration (Fig. S2A). The binding curve yielded a [Mg^2+^]1/2 of ∼14 mM and is consistent with 3 bound Mg^2+^ ions. Since deprotonation of the 3′ OH is an important step in ligation, we measured *k*obs at pH values between 6 and 10 (Fig. S2B). Ligation rates increased between pH 6 and pH 9 and then fell, as in the corresponding uncatalyzed reaction.(47) The ‘log *k*obs vs. pH’ curve is linear between pH 6 and pH 9 in both CS3-catalyzed and uncatalyzed PPP-ligation. While a slope of 1, as observed for the uncatalyzed reaction likely indicates 3′ OH deprotonation in the rate-determining step, the slope of 0.45 for the CS3-catalyzed ligation may reflect a change in mechanism. Similar deviations were observed for the parent AIP-ligase, RS1,(9) a DNAzyme that ligates imidazole-activated RNA,(57) and for variants of the class I ligase.(58)

The catalytic promiscuity of CS3 as evidenced by its ability to catalyze ligation with both PPP-Lig and AIP-Lig was surprising as its parent, RS1 is specific to 5′-monophosphorylated RNA oligonucleotides activated with 2-aminoimidazole and does not ligate triphosphorylated RNA.(9) The specificity of RS1 is underscored by its lack of reactivity toward a substrate activated with 2-methylimidazole (MeIP), which suggests that the RS1 ribozyme may interact with the amino group of 2-aminoimidazole. In contrast, CS3 ligates MeIP-Lig, albeit less efficiently than AIP-Lig (Fig. S3). This observation is consistent with RS3 having a flexible or open active site that accommodates diverse leaving groups such as pyrophosphate, 2-aminoimidazole, and 2-methylimidazole.

### Acquisition of a new function is accompanied by the emergence of a new structure

The SHAPE-derived secondary structure of the catalytic domain of the parent AIP-ligase, RS1, consists of two stems interrupted by an internal loop with the outer stem closed by a hairpin loop (Fig. S4A). The inner stem, which was part of the constant region in the library, is flanked on its 5′ side by a 25 nt sequence that was designed as a PCR primer binding site and a U6 linker connected to the 8 nt primer sequence on its 3′ side.(9) SHAPE probing of CS3 revealed a new secondary structure, which is composed of two stems interrupted by internal bulges and terminating in stem loops (Fig. S5). These stems are connected by a stretch of unpaired nucleotides, which could potentially act as a hinge between these two stem regions. The variable region (shown in orange in Fig. 3A) extensively base pairs with the fixed 5′ primer binding region and the nucleotides that formed the 3′ strand of the inner stem in the RS1 structure (Figs. 2A, 3A). The loss of activity upon 5′ truncation (Fig. 3C) and a high degree of nucleotide conservation in the variable region (Fig. S6) are both consistent with this secondary structure. A stretch of 7 nucleotides (5′-GCGGAAU-3′) connecting the two stem regions is perfectly complementary to part of the substrate that remains unpaired upon template binding, referred to here as the 3′ overhang (5′-AUUCCGC-3′) (Fig. S4B). While some of these nucleotides are free to pair with the substrate (complementarity is indicated by black dashed lines), nucleotides 3 to 5 (5′-GGA-3′)) may be base paired within the ribozyme secondary structure (complementarity with the substrate is indicated by magenta dashed lines). Disrupting the putative base-pairing between CS3 and the substrate by sequestering the substrate 3′ overhang with longer templates eliminated ligation (Fig. S4B, D). Similar results were obtained with RS1, where nucleotides in its stem loop (nucleotides 16-21, 23) were complementary with the substrate 3′ overhang (Fig. S4A, C). These results suggest that both CS3 and RS1 use base-pairing with the substrate to assemble the enzyme-substrate complex. Despite having distinct sequences and structures, the two ligases have converged on a similar solution to substrate binding.

The structural differences between RS1 and CS3 were further highlighted by a combination of SAXS analysis and molecular dynamics simulation (Fig. S7). First, we used SAXS to generate low resolution molecular envelopes for RS1 and CS3. Next, we used Rosetta’s FARFAR2(59) method to generate molecular models for RS1 and CS3 with the corresponding SHAPE-derived secondary structures included as constraints. These computational models were used as starting points for all-atom molecular dynamics simulations. Computed molecular models of RS1 and CS3 were fitted to experimental SAXS envelopes (Fig. S7A-F). While the best computed model of RS1 agreed with the SAXS envelope, the best fit of CS3 to the SAXS data required a two-state model (Fig.S7 C, F). The structural ensembles qualitatively suggest a more dynamic structure for CS3, which agrees with the two-state fit to the SAXS data. A higher degree of flexibility in the CS3 structure is also indicated by a greater root mean square deviation (RMSD) value of the CS3 structure during the simulation with respect to the starting conformation (Fig.S7 G, H).

### Population dynamics of selected ligases

The parent AIP-ligase ribozyme, RS1, underwent drastic changes to its catalytic scaffold to adapt to PPP-ligation (in CS3). This is a direct consequence of the large distance of 28 mutations between RS1 and CS3, which represents 70% sequence change within the 40 nt variable region, and an overall change of ∼30% (28 nt/95 nt). This large mutational distance was unexpected considering that the level of mutagenesis in the starting library was only 21%. The population of RS1 variants moved farther from RS1 along the selection trajectory, as visualized by high-throughput sequencing (Fig. 5A). Sequences 8 to 11 mutations from RS1 were the most abundant after the round 1, but the maximum of this distribution shifted to variants with 10 to 13 mutations after round 3. After round 4, concurrent with detectable PPP-ligase activity, the selected RNA population became dominated by three distinct populations that were 13, 16, and 28 mutations from RS1. Sequences 13 and 16 mutations from RS1 represent unrelated ligase clusters CS1 and CS2 + CS5, respectively. Sequences 28 mutations from RS1 represent the PPP-ligase cluster, CS3. Although CS3 and RS1 catalyze similar ligation reactions, CS3 is mutationally farther from RS1 than the unrelated ligases. The isolation of variants that differ significantly from the parent sequence to access related functions has been observed consistently during *in vitro* evolution experiments.(28–33)

To study the population dynamics of sequences within the CS3 cluster, we plotted the fractional abundance of all CS3 variants with >100 reads for each round (Fig. 5B, Table S2). Most sequences become more abundant during selection, with a sudden enrichment after round 4. While the peak sequence, CS3 (sequence 1 in Fig. 5B), remained dominant in the cluster throughout, ultimately populating >50% of the cluster, certain sequences such as 22, 26, 48, and 56 became gradually more abundant after emerging in round 2 (i.e., with >100 reads). On the other hand, the abundance of sequences 53, 55, 56, and 62 fell below the read threshold but then re-emerged later during the selection. Tracking the relative abundances of PPP-ligase variants illuminate how these ribozyme sequences respond to changing selection pressures in each round.

**Fig. 5.**
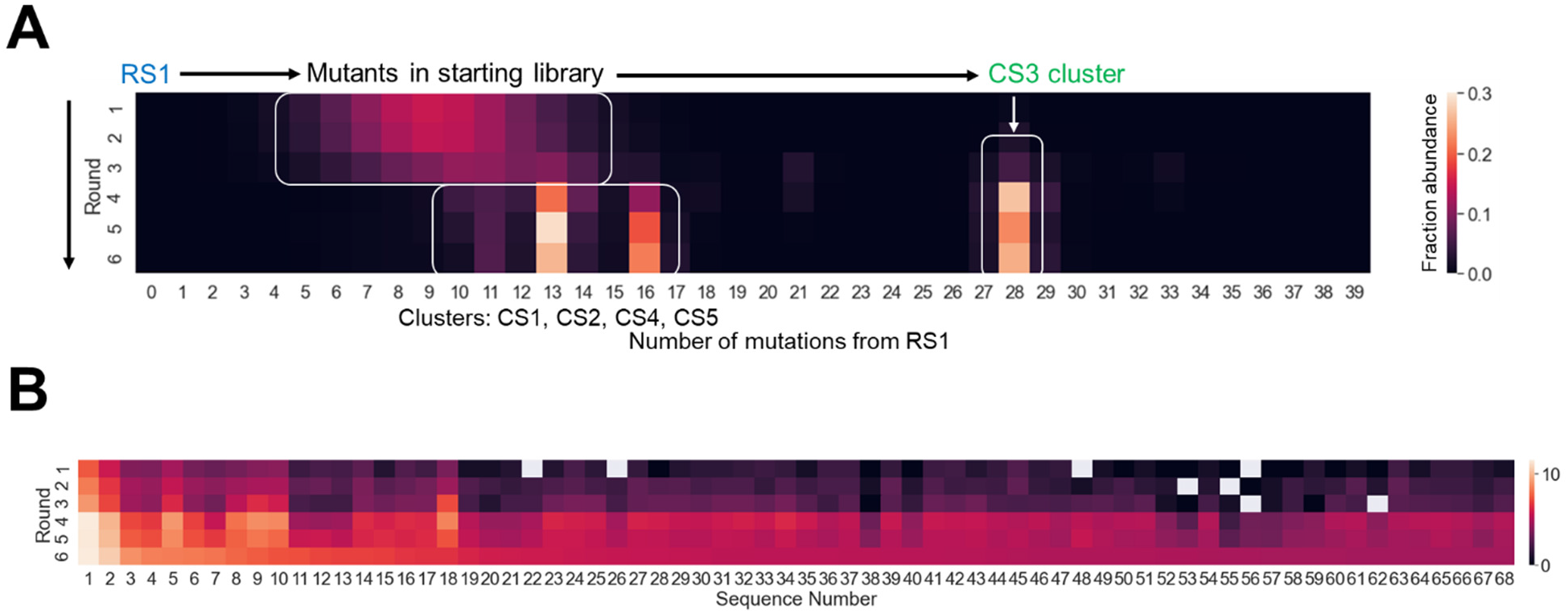
RNA population dynamics during *in vitro* evolution. **A.** The selected sequence population moved farther from the parent sequence, RS1, along the selection trajectory forming distinct clusters. While ligases CS1, CS2, CS4, and CS5 and their close variants were 10-16 mutations away from RS1, the PPP-ligase, CS3, was 28 mutations from RS1. Colors indicate fractional abundance in the entire sequenced pool. Only sequences that have >100 reads are shown. **B.** Sequence dynamics in the CS3 cluster. The peak sequence, CS3 (Sequence 1), was already selected in round 1 and became more abundant during selection. Certain sequences like Sequences 53, 55, 56 and 62 fell below the read threshold but re-emerged in later rounds. See Table S2 for information about the sequences and their relative abundance. Colors indicate log of read counts within the CS3 cluster. Only sequences that have >100 reads are shown.

### A quasi-neutral pathway facilitates evolutionary innovation

The large mutational distance between the AIP-ligase, RS1 and the PPP-ligase, CS3 raises questions about the feasibility of adaptive mutational walks along the ligase fitness landscape. It has been proposed that the RNA sequence space is permeated by neutral networks in which each mutant intermediate is one mutation from its immediate neighbors and retains fitness.(41, 43, 60) The existence of neutral pathways connecting distinct RNAs drastically reduces the need for mutations that significantly increase fitness (adaptative mutations). While our results show that a PPP-ligase can be evolved from an AIP-ligase in a single jump following intensive mutagenesis, we wondered if these ligases are connected by neutral pathways, where each single-step mutant intermediate retains at least some ligase function. To identify these neutral intermediates, we computationally generated all possible single mutants of CS3 and ran Dijkstra’s shortest path finding algorithm from RS1 to these single mutants, using only the sequences present in the sequencing data obtained from rounds 1-6. The algorithm was designed to identify the path that terminated in a sequence that most closely resembles CS3. Unfortunately, the path terminated in a sequence that was only 2 mutations from RS1 (INT3 in Table S3), suggesting the absence of intermediates between RS1 and CS3 in the sequencing data. Next, we repeated this step by running our algorithm from CS3 to all single mutants of INT3. This search terminated at INT4 (Table S3), which is 24 mutations from INT3 and 3 mutations from CS3. Collectively, this computational approach revealed six intermediate sequences present in the sequencing data – three closely-resembling RS1 (INT1-INT3) and three closely-resembling CS3 (INT4-INT6), with a 24-mutation gap between INT3 and INT4 (Table S3). This gap does not necessarily indicate the absence of a neutral path as our experiment does not sample the entire sequence space between RS1 and CS3. Therefore, we took an experimental trial and error approach to fill this gap by designing and assaying the AIP and PPP ligase activities of mutant sequences that complete a single-step mutational pathway between INT3 and INT4. Only mutant sequences with rate enhancements that were >10% of that exhibited by target sequence, CS3 were considered as ‘quasi-neutral’ intermediates and included in our analysis (Fig. 6, Table S4). As CS3-catalyzed AIP ligation and PPP ligation are ∼100- and ∼1000-fold faster than their background reactions, respectively, intermediates had to show rate enhancements of 10-fold for AIP-ligation and 100-fold for PPP-ligation to be considered quasi-neutral.

INT4 (mutant 26 in Fig. 6), which represents the mutational jump from the AIP-ligase phenotype to the PPP-ligase phenotype, catalyzed both AIP and PPP-ligation as expected by its sequence similarity to CS3. The two intermediates between INT4 and CS3 were also active for both functions. We made sequential mutations to RS1 with the aim of delineating a neutral pathway to INT4 that passes through the intermediate sequences identified by our path-finding algorithm (INT1, INT2, and INT3). While the first three mutations collectively lowered AIP-ligase activity by only ∼4 fold, the fourth mutation caused a marked reduction. The subsequent mutations generated sequences that exhibited lower levels of AIP-ligase activity, including some sequences that showed activities just above the 10% threshold. The lower activities of these intermediates may result from the adoption of an ensemble of RNA folds, most of which are inactive, as observed for ‘neutral’ intermediates between the class III ligase and HDV ribozymes.(61) PPP-ligase function emerged after 26 mutations, at which point there was also a significant rise in AIP-ligase activity. This could be due to the creation of a stable, catalytic structure resembling CS3. All intermediates that exhibit PPP-ligation on this path are part of the CS3 cluster. The appearance of PPP-ligase activity only in mutants that are close to CS3 suggests that CS3 may be far from being optimal for PPP ligation; if so, further exploration of the sequence space surrounding CS3 may reveal more active sequences. As the new CS3-like fold also catalyzed AIP-ligation, CS3 represents a distinct solution to AIP-ligation, in addition to functioning as a PPP-ligase. Our results also show that although RS1 is 28 mutations from CS3, sequences with exclusive AIP-ligase activity may be found in close proximity to PPP-ligases.

To understand how the evolution of RNA structure correlates with the evolution of catalytic function, we used computational structure prediction to follow the structural changes along the quasi-neutral mutational path from RS1 to CS3. We found that new intermediate structures were formed along the mutational path through the formation or disruption of single base pairs, one at a time. The structures could be grouped into seven distinct structure-types that include an RS1-like structure type, a CS3-like structure-type, and five new structure types connecting these two (Fig. S8). While structural changes are more-or-less gradual with a notable absence of base-pairing between nucleotides of the constant (black) and variable (blue and green) regions in most structures (Fig. S8A-W), we observed a sudden and significant change in the overall fold after the 23^rd^ mutation. The CS3-like fold of Int23, which emerged suddenly due to a single mutation to Int22, is created by extensive base-pairing between nucleotides from the constant and variable regions (Fig. S8X). This can be considered a molecular version of punctuated equilibrium, often seen in experimental RNA evolution.(42, 60) Taken together, our results provide experimental support for computational studies in the 1990s that emphasized the importance of multiple genotypes mapping on to a single structural phenotype in allowing neutral pathways to connect distinct structural folds (and functions) in RNA.(42, 62) Although evolution operates on RNA sequences, selection pressures operate in the RNA structure space. Therefore, RS1 and CS3, two distinct RNAs, may be far in sequence space, but they need to traverse just five distinct structures to interconvert. Moreover, the existence of multiple structural phenotypes for a single functional phenotype (e.g., AIP ligation) underscores the importance of preserving fitness in the face of mutations, i.e., robustness. Robustness has been proposed to be critical for evolution as it allows mutational excursions on the fitness landscape and consequently, chance encounters with new functions. The mutational path between RS1 and CS3, outlined in this work supports the crucial role of robustness in enhancing the evolvability of a system.(39, 40)

**Fig. 6.**
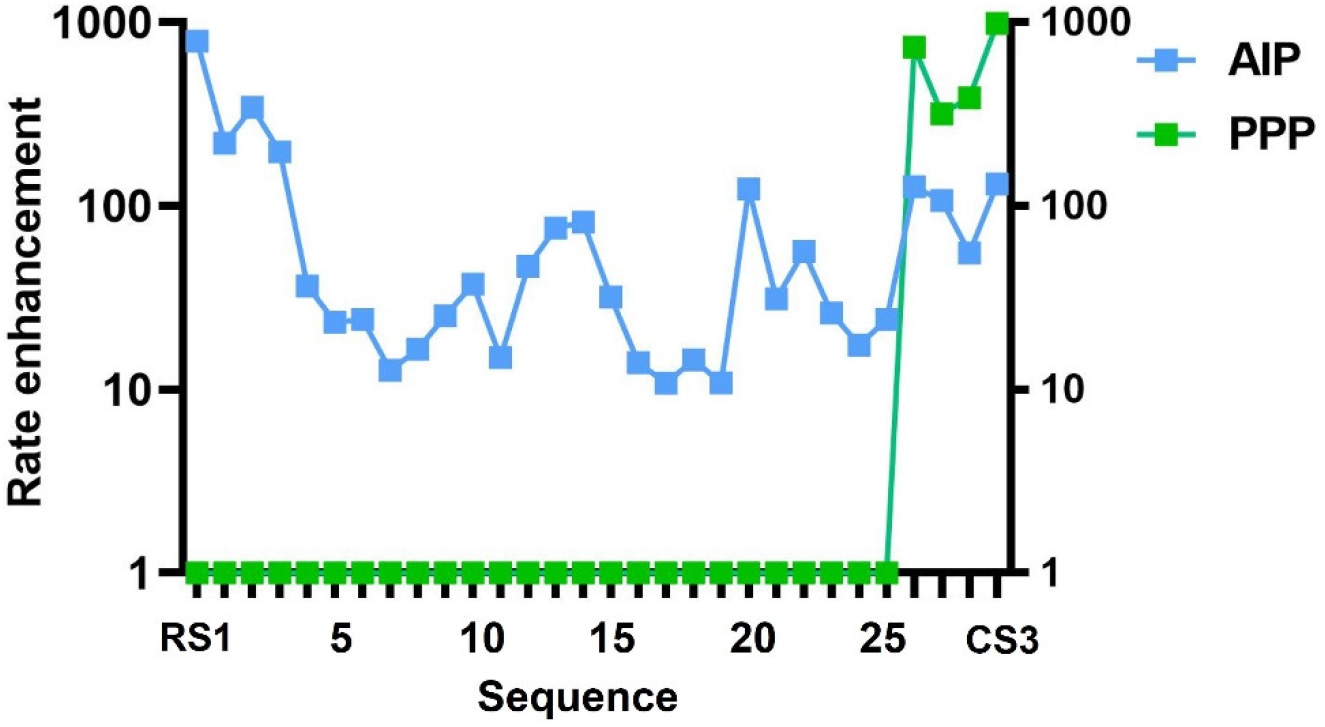
A quasi-neutral mutational pathway connecting RS1 and CS3. AIP-ligation and PPP-ligation are depicted in blue and green, respectively. Each sequence, represented by a data point, is separated from its immediate neighbors by a single mutation (i.e., Hamming distance of one). An accumulation of point mutations to RS1 causes a fall in AIP-ligase activity, which fluctuates until PPP-ligase activity emerges in Sequence 26 (INT4 in Table S3), which is also accompanied by an increase in AIP-ligase activity. This represents the emergence of a new stable but promiscuous catalytic fold. Sequences 1-4 are predicted to adopt an RS1-like structure, sequences 23-28 are predicted to adopt a CS3-like structure, and sequences 5-22 are predicted to fold into structures that can be grouped into five distinct structure-types (for secondary structures, see Fig. S8). Sequences used in this analysis are listed in Table S4. Ligation reactions contained 1 µM ribozyme, 1.2 µM RNA template, and 2 µM RNA substrate (AIP-Lig or PPP-Lig) in 100 mM Tris-HCl pH 8.0, 300 mM NaCl, and either 10 mM MgCl2 (AIP-Lig) or 100 mM MgCl2 (PPP-Lig).

## DISCUSSION

We have demonstrated that ribozymes that catalyze RNA ligation using prebiotic phosphorimidazolide substrates (AIP-ligase) may be evolved to ligase ribozymes that utilize triphosphate substrates (PPP-ligase). The ability of an AIP-ligase ribozyme to morph into a very different structure with the ability to ligate both PPP- and AIP-substrates is consistent with the possibility that a primitive state of RNA-catalyzed RNA assembly using prebiotically-relevant phosphorimidazolide substrates transitioned to an intermediate stage where ribozymes used triphosphate substrates for RNA assembly, thus setting the stage for the later transition to modern protein-catalyzed RNA synthesis using triphosphate substrates. The PPP-ligase sequence reported here adds to the list of ligase ribozymes that catalyze PPP-ligation. Our new PPP-ligase is distinct from most previous examples in that it catalyzes PPP-ligation on an external template, which would be critical for a ribozyme to be able to carry out general RNA assembly processes. This is in contrast to, for example, the class I ligase, where the template is supplied by a region in the ribozyme itself and the triphosphate group is on the ribozyme 5′ terminus, which reacts with a 2′/3′-hydroxyl group of the ‘substrate’.(19)

A remarkable feature of the evolution of the new PPP-ligase from the prior AIP-ligase is the dramatic shift in the folded structure of the ribozyme. Given that the RS1- and CS3-catalyzed ligation reactions are quite similar except for their leaving groups (2-aminoimidazole vs. pyrophosphate), one might have expected this change to require only a minor readjustment of the ribozyme structure. In reality, it appears that access to the new PPP-ligase function required a change to a radically different RNA fold. The creation of new folds to perform apparently similar functions, as opposed to minor adjustments to a binding site or active site, has been observed during previous examples of the directed evolution of aptamers and ribozymes,(28–33) consistent with the idea that RNA fitness landscapes are rugged, such that smooth structural morphing between different functional states is not generally possible. We were surprised to find that the new PPP-ligase is promiscuous, in that it can catalyze ligation with AIP- and PPP-substrates, even though it was selected only for PPP-ligation. Why directed evolution resulted in a new structure that supports both the new, selected, function, while also retaining the old function is unclear. Active site promiscuity has been observed previously in functional RNAs found in nature and evolved artificially, so one possibility is that ribozyme active sites are inherently likely to be non-specific and therefore promiscuous.(63–67) An alternative possibility is that the observed promiscuity might arise from the ability of the CS3 sequence to adopt two distinct catalytic folds corresponding to PPP-and AIP-ligation. However, CS3 cannot fold into the parental RS1-like structure, so the dual functionality cannot result from an ability to fold into both the old and new structures. If the AIP-ligase activity is due to a distinct folded structure, that structure would have to have acquired AIP-ligase activity in the absence of selection, which seems unlikely. Furthermore, the only two ribozyme sequences known to populate two distinct catalytic structures were designed by extensive sequence engineering,(43, 44) making it unlikely that CS3’s promiscuity is derived from structural plasticity. We therefore favor a model in which the active site of the CS3 ribozyme is itself less specific, perhaps through an ability to bind both the 2AI and the pyrophosphate leaving groups. Regardless of the underlying mechanism, the promiscuity of the PPP-ligase makes it an interesting example of a potential intermediate in the transition from nonenzymatic RNA assembly to RNA assembly with more biologically relevant substrates. Ribozyme promiscuity could have been an important driver of evolutionary innovations in an RNA World by providing opportunities for divergent functions to arise within one sequence, with subsequent evolutionary optimization following gene duplication events.(38, 43, 68, 69)

We outlined a mutational path that connects the parental RS1 ligase with the evolved CS3 ligase, where each intermediate differs from its predecessor by a single mutation, and where all intermediate sequences retain at least some ligase activity. This is the third example where two distinct catalytic RNA folds are connected by a quasi-neutral path.(43, 44) The existence of such a neutral path implies both the existence of a very large number of sequences that support a single function (robustness) as well as a smaller number of sequences that support both functions (promiscuity). Due to the high dimensionality of the RNA sequence space, the quasi-neutral path we identified between RS1 and CS3 is likely just one of many neutral paths that collectively constitute a neutral network bridging these two catalytic functions. In the neutral pathway outlined in this work, the sequence intermediate between the source sequence (RS1) and target sequence (CS3) show lower activities than RS1 and CS3, which would make a step-wise transition between these two functions difficult in a biological system with low mutation rates. Mutations that disrupt the overall fold or the catalytic apparatus of RS1 likely explain the reduced activities of these intermediate sequences. It is possible that neutral networks with more active intermediate states may be revealed through additional high-throughput sequencing and computational approaches.(37, 70)

One puzzling feature of our selection experiment is that the novel CS3 ligase differ by many mutations (28 out of 40 mutagenized sites) from the parental sequence. Sequences with 28 or more mutations constituted only 4 x 10^-11^ of the initial doped library, corresponding to roughly 4 x 10^4^ sequences. Based on previous selection experiments, it would be extremely unlikely for a novel ribozyme structure to emerge from such a small pool of sequences. Even if we assume that all 28 mutations were not essential, so that the new ribozyme could have emerged from the pool of sequences with, for example, 25 or more mutations, the available pool would still correspond to less than 2 x 10^7^ sequences, whereas most novel ribozymes have emerged in previous selection experiments with a frequency of less than 10^-10^ random sequences. A possible explanation for this apparent discrepancy is that the new ribozyme first emerged from a less active sequence with fewer mutations that was present in the original doped library. Additional mutations, conferring enhanced activity, could have arisen during the amplification stages of the first few rounds of selection. However, these hypothesized ancestral sequences must be extremely rare, such that we did not detect them by deep sequencing.

Ribozyme-catalyzed RNA assembly using triphosphate building blocks would have been evolutionarily advantageous even in the early phases of the RNA World due to the greater stability of PPP-substrates to hydrolysis compared to AIP-substrates. As RNA assembly reactions with PPP-substrates yield pyrophosphate as a side-product, a self-sustaining cycle may have arisen in which phosphorimidazolide substrates were converted to triphosphate substrates by attack of pyrophosphate. Such a cycle may have facilitated the transition from prebiotic phosphorimidazolides to more biotic triphosphates as preferred substrates for enzyme-catalyzed RNA assembly. Since biological RNA assembly occurs via the polymerization of triphosphorylated monomers, and not via oligomer ligation, we speculate that PPP-ligase ribozymes may have evolved to utilize NTPs. Ribozyme catalyzed RNA synthesis from NTPs may have thereby set the stage for protein polymerases to co-opt NTPs as substrates for RNA assembly. Alternatively, the first protein enzymes to catalyze RNA assembly could have been ligases that used triphosphorylated oligomers as substrates. Ligases are expected to be simpler catalysts than polymerases as they do not need to bind monomeric substrates. Instead, oligonucleotide substrates can use base-pairing to bind to RNA templates. Furthermore, ligase enzymes do not need to be as active as polymerases, since fewer phosphodiester bond forming steps are required to assemble an RNA of any given length. Consequently, ligases are expected to be more abundant in both RNA and protein sequence space and are likely to have played an important role in the early stages of modern biology.(71) Indeed, protein enzymes that catalyze RNA ligation using triphosphorylated RNA oligonucleotides have been identified using laboratory evolution.(72) Regardless of the exact biochemical events that led to the emergence of modern biology, our work provides an experimental model that connects the past and present of RNA assembly.

## MATERIALS AND METHODS

### Materials

All chemical reagents were purchased from Sigma, unless otherwise specified. The hydrochloride salt of 1-ethyl-3-(3-dimethylaminopropyl) carbodiimide was purchased from Alfa Aesar. The hydrochloride salt of 2-aminoimidazole was purchased from Combi-Blocks, Inc. Enzymes were purchased from New England Biolabs unless mentioned. SYBR Gold Nucleic Acid Gel Stain was purchased from ThermoFisher Scientific. 100% ethanol was purchased from Decon Laboratories, Inc. QIAquick PCR purification kits were purchased from Qiagen. The Sequagel-UreaGel concentrate and diluent system for denaturing polyacrylamide gels was purchased from National Diagnostics. Oligonucleotides used in this work are listed in Supplementary Table S6 and were purchased from Integrated DNA Technologies (IDT) except for PPP-Lig and PPP-LigB, which were purchased from Chemgenes.

### RNA preparation and substrate activation

Ribozyme constructs were prepared by *in vitro* transcription of PCR-generated dsDNA templates containing 2′-*O*-methyl modifications at the last two nucleotides of the template strand to reduce transcriptional heterogeneity at the 3′ end of the RNA.(73) Each 1 ml transcription reaction contained 40 mM Tris-HCl (pH 8), 2 mM spermidine, 10 mM NaCl, 25 mM MgCl2, 10 mM dithiothreitol (DTT), 30 U/mL RNase inhibitor murine, 2.5 U/mL thermostable inorganic pyrophosphatase (TIPPase), 4 mM of each NTP, 30 pmol/mL DNA template, and 1U/µL T7 RNA Polymerase and were incubated at 37°C for 3 h. DNA template was DNase I-digested (5U/mL at 37° C for 30 min) and the RNA was extracted with phenol-chloroform-isoamyl alcohol (Invitrogen), ethanol precipitated, and purified by 10% denaturing PAGE. Ligation templates, FAM-labeled primers, and ssDNA were purchased from Integrated DNA Technologies.

CS3 variants with modified 5′ chemistries (5′P and 5′OH) were generated by the splinted ligation of three RNA oligonucleotide pieces (Table S6). The first piece contained either a 5′P or a 5′OH modification. The second and third pieces were 5′-monophosphorylated to enable ligation by T4 RNA ligase 2. 60 pmol of each piece were incubated with 40 pmol each of the two DNA splints at 90 °C for 3 min followed by a 10 min incubation at 30 °C. This was followed by the addition of 1U/ µL RNA ligase 2 and 1X T4 RNA ligase buffer. The 20 µL reaction was incubated at 30 °C for 2 h, the RNA was extracted with phenol-chloroform-isoamyl alcohol, and purified by 10% denaturing PAGE.

The 5′-monophosphorylated oligonucleotide corresponding to the substrate sequence (Table S6) was activated by as previously described.(9) Briefly with 0.2 M 1-ethyl-3-(3 dimethylaminopropyl) carbodiimide (HCl salt) and 0.6 M 2-aminoimidazole (HCl salt, pH adjusted to 6) in aqueous solution for 2 h at room temperature, followed by four or five washes with 200 µL water per wash in Amicon Ultra spin columns (3kDa cutoff).This was followed by reverse phase analytical HPLC purification using a gradient of 98% to 75% 20 mM TEAB (triethylamine bicarbonate, pH 8) versus acetonitrile over 40 min.

### *In vitro* evolution of triphosphate ligase ribozymes

The selection library was derived from the most active AIP-ligase (called RS1) identified from an earlier selection.(9) The 40 nt region in RS1 that constituted the library region in that selection was subjected to mutagenesis, where each nucleotide was doped at 21%. At each position the probability of retaining the parental nucleotide is 79%, and the other three nucleotides occurred with a 7% probability, each. The 21%-doped ssDNA library was purchased from IDT. 1.2 nmol ssDNA was converted to dsDNA by reverse transcription. The ssDNA was annealed with 1.2 nmol PCR_Rvs_primer (Table S6) and incubated with 2 mM dNTP mix, 2 mM DTT, 1X Protoscript buffer, and 5U/ μL ProtoScript II reverse transcriptase at 42 °C for 4 h. The reaction products were purified using QIAquick PCR purification kit. The RNA library used for the first round of selection was obtained by transcribing the dsDNA as described above.

The selection strategy for isolating ribozymes capable of 5′-triphosphate RNA ligation involved partitioning active from inactive sequences by streptavidin bead capture. The partially randomized RNA library (1 µM) was incubated at room temperature with an RNA template (1.1 µM), 100 mM Tris-HCl, pH 8, 250 mM NaCl, and substrate, PPP-LigB (1.2 µM). The amount of RNA library used in each round, the incubation times, and the Mg^2+^ concentrations for each round are listed in Table S1. Reactions were quenched by adding 200 mM EDTA, 200 µM Quench_Oligo1 (a deoxyoligonucleotide complementary to the substrate) and 20 µM Quench_Oligo2 (a deoxyoligonucleotide complementary to the 3′ end of the ribozyme-primer construct) (Table S6). The reaction mixture was heated to 95 °C for 2 min, cooled to RT, then incubated with MyOne Streptavidin C1 Dynabeads (1 mg/100 pmol RNA). Before use, Dynabeads were prepared by washing twice with buffer 1 (1 M NaCl, 1 mM EDTA, 10 mM Tris pH 7.6, 0.2% Tween-20), once with buffer 3 (25 mM NaOH, 50 mM NaCl, 1 mM EDTA), then twice more with buffer 1. 1 mL buffer was used for washing 1 mg beads. Beads were blocked to reduce non-specific RNA retention by tumbling in 1 mL buffer 1 containing 10 μg/mL tRNAs for 1 h. Following blocking, beads were washed twice with buffer 1, and the quenched reaction mixture was added to the beads in the presence of buffer 1. To immobilize biotinylated RNAs, the solution was tumbled for 30 min. The beads were washed with buffer 1 twice, then tumbled with buffer 2 (8 M urea, 1 M NaCl, 1 mM EDTA, 10 mM Tris pH 7.6, 0.1% Tween-20) for 20 min, twice, then tumbled with buffer 3 for 5 min, twice. After three washes with buffer A and removal of liquids, 100 μL of the elution mix (95% formamide, 10 mM EDTA) was added to the beads and the beads were incubated at 65 °C for 6 min to elute the immobilized RNA from the beads. The eluted RNA was ethanol precipitated by adding 200 µL water, 50 µL sodium acetate (pH 5.2), 2 µL glycogen (20 mg/ml; RNA grade - ThermoFisher Scientific) and 900 µL 100% cold ethanol and storing at -80°C for 30 min. The pellet was washed with 70% ethanol and air dried. All washes were with 1 mL buffer and a Dynal magnetic tube rack was used for magnetic separation.

The pellet was resuspended in water and reverse transcribed with ProtoScript II reverse transcriptase and reverse transcription primer, RT_primer (Table S6) at 42 °C for 3 h as described before. The enzyme was heat inactivated at 80 °C for 5 min and the product was purified with QIAquick PCR purification kit. The purified product was directly used as input for a PCR with primers, PCR_Fwd_primer and PCR_Rvs_primer (Table S6). The PCR cycle was: 1) 95 °C for 2 min; 2) 10 cycles of 94 °C for 30 s, 54 °C for 1 min., and 72 °C for 1 min.; 3) 72 °C for 10 min. The PCR product was purified using QIAquick PCR purification kit and 100 ng of the PCR product was further amplified in a second PCR as above for 8 cycles. These steps were taken to minimize amplification due to the presence of RT_primer, which may complete with PCR_Rvs_primer during PCR. The dsDNA from this PCR was transcribed to generate the input RNA library for the next round and was also subjected to the three sequencing PCRs to generate the corresponding sequencing library for high-throughput sequencing (see ‘Library preparation for high throughput sequencing’).

### Ligation assays

Ligation reactions contained 1 µM ribozyme, 1.2 µM RNA template, and 2 µM RNA substrate in 100 mM Tris-HCl pH 8.0, 300 mM NaCl, and either 10 mM (AIP-Lig) or 100 mM (PPP-Lig/PPP-LigB) MgCl2. Aliquots were quenched with 5 volumes of quench buffer (8 M urea, 100 mM Tris-Cl, 100 mM boric acid, 100 mM EDTA) and analyzed on a 10% denaturing PAGE. Gels were stained using SYBR Gold and imaged on an Amersham Typhoon RGB instrument (GE Healthcare), Scans were analyzed in ImageQuant IQTL 8.1. Kinetic data were nonlinearly fitted to the modified first order rate equation, y = A (1 – e-^kx^), where A represents the fraction of active complex, k is the first order rate constant, x is time, and y is the fraction of ligated product in GraphPad Prism 9.

### Library preparation for high throughput sequencing

RNA pools were sequenced after every round of selection. To prepare the selected pools for sequencing, sequencing adaptors were introduced via three rounds of PCR with different primer sets (Table S6), using the dsDNAs from each pool that were used to generate the RNA pool for the next round. In the first round, 25 ng dsDNA was amplified in a 100 μL PCR containing 0.5 μM primers, SeqPCR_primer1 and SeqPCR_primer2, by Taq DNA polymerase for 4 cycles. The PCR product was purified using QIAquick PCR Purification Kit (Qiagen), and then further amplified for 7 cycles in the second round of PCR containing 40 ng dsDNA, 0.5 μM primers, SeqPCR_primer3 and SeqPCR_primer4. The purified product of the second PCR was amplified in a 50 μL reaction containing 1 unit of Q5 Hot Start High-Fidelity DNA Polymerase, 100 ng of ds DNA, 1X Q5 buffer, 0.2 μM each of SR Primer for illumina and Index Primer for illumina (New England Biolabs), and 80 μM dNTPs. This third round PCR mixture was incubated at 98 °C for 30 s, followed by 6 cycles of 98 °C for 10 s, 62 °C for 15 s, and 72 °C for 20 s, and finally, 72 °C for 2 min. The PCR product was purified using QIAquick PCR Purification Kit and then on a 1.4% agarose gel. Purified DNAs were extracted from agarose gels by Freeze N’ Squeeze gel extraction spin columns (Bio-Rad). The DNA was purified from the residual dye and concentrated using an Oligo Clean & Concentrator Kit (Zymo Research) and sequenced in an illumina MiSeq instrument.

### Bioinformatics analysis of high throughput sequencing data

Sequencing reads from each round were pre-processed using Python scripts as follows. Sequences were first filtered for the presence of the predefined eight base-pair constant stem region and only sequences that passed this filter were subject to quality control. Sequences that satisfied the quality score threshold of 20 for each nucleotide position (Q ≥ 20) with less than one percent error were trimmed to the 40 nt variable region.(74) Trimmed sequences were dereplicated using a custom script and clustered using the Clustal Omega algorithm.(75, 76)

The relative abundances of CS1-CS5, the peak sequences of the five most abundant clusters, were plotted across rounds to generate Fig. 2C. To generate Fig. 5A, we counted the number of reads for sequences at different mutational distances from the parent AIP-ligase, RS1, and represented their relative abundances for rounds 1-6 in a heatmap. Only sequences with ≥100 reads are shown. Similarly, Fig. 5B was generated by plotting the fractional abundances of sequences (with ≥100 reads) in the CS3 cluster across rounds 1-6. To generate Fig. S6A, we counted the frequency with which each of the four nucleotides occur at a given position in the CS3 peak sequence and plotted the probability fraction against nucleotide number. This reveals nucleotide conservation within the CS3 cluster. Fig. S6B was generated using the WebLogo online server (https://weblogo.berkeley.edu/logo.cgi).

### Regiospecificity of the phosphodiester bond between CS3 and PPP-Lig

∼4 pmol of the purified ligated product obtained from an overnight ligation reaction between CS3 and PPP-Lig was digested with 20 U of RNase R (Lucigen) in a 10 μL reaction in the presence of 1X RNase R buffer at 50 °C for 1 h. The reaction was quenched by adding 0.3 μL of 0.5 M EDTA and the enzyme was deactivated by heating at 95 °C for 3 min. 10 μL of quench buffer (8 M urea, 100 mM Tris-Cl, 100 mM boric acid, 100 mM EDTA) was added to the heat-inactivated reaction, and the reaction was analyzed by 10% denaturing PAGE.

### Secondary structure determination by SHAPE

SHAPE probing of CS3 was performed using a protocol reported in Walton *et al.*, 2020.(9) The sequence used for SHAPE probing (CS3_SHAPE) consisting of 5′ and 3′ SHAPE cassettes is included in Table S6. Briefly, 100 pmol CS3_SHAPE was folded in 100 mM Tris-HCl, pH 8, 250 mM NaCl, 10 mM Mg^2+^ and divided into ‘modification’ and ‘control’ tubes. SHAPE reagent, 1M7 (in DMSO) was added to a final concentration of ∼40 mM in the ‘modification’ tube and only DMSO was added to the ‘control’ tube. Both modified and unmodified RNAs were reverse transcribed using 40 pmol of a 5′ FAM-labeled primer (SHAPE_RT_primer) and Superscript III reverse transcriptase (Invitrogen) (Table S6). Reverse transcription of 30 pmol unmodified RNAs in the presence of each of the four ddNTPs and 25 pmol SHAPE_RT_primer was used to generate sequencing lanes. Reactions and quenched and analyzed by 10% denaturing PAGE. Normalized SHAPE reactivity was calculated by first excluding the most reactive nucleotide position and then dividing reactivities at each position by the average of the 10% most reactive positions. Normalized SHAPE reactivities were used to constrain secondary structure prediction in the RNAstructure program.(77)

### Small-angle X-ray scattering analysis

SAXS data were collected at the SIBYLS beamline at the Advanced Light Source (Berkeley, CA) as previously described.(78–80) For CS3, two concentrations were analyzed: 0.5 mg ml^-1^ and 1 mg ml^-1^. For RS1, three concentrations were analyzed: 0.5, 1, and 2 mg ml^-1^. Data were collected in 10 mM MgCl2, 200 mM NaCl, and 200 mM Tris pH 7.5 at 20°C. The X-ray wavelength was set to 1.127 Å with a sample-to-detector distance of 2,100 mm, which gives scattering vectors (q) ranging from 0.01 Å-1 to 0.4 Å^-1^. The SAXS flow cell was coupled to an Agilent 1260 Infinity HPLC system using a Shodex 802.5 SEC column equilibrated with the running buffer as indicated above with a flow rate of 0.65 ml min^-1^. For SAXS measurements, 2 second X-ray exposures were collected during a 25-minute elution. Buffer subtraction was performed with buffer matched controls. For CS3, the 0.5 mg ml^-1^ sample provided the best Guinier fit and was used for subsequent analysis. For RS1, the 1 mg ml^-1^ sample provided the best Guinier fit and was used for subsequent analysis.

The radius of gyration (Rg) was calculated for each buffer subtracted frame using the Guinier approximation in the program RAW.(81) Final merged SAXS profiles obtained by integration of multiple frames at the elution peak were used for further analyses. The volume of correlation (Vc) was used to estimate molecular weight,(82) and the pair distribution function [P(r)] was used to calculate maximum inter-particle distances (Table S5).(83) Molecular envelopes for each ribozyme were calculated using DENSS(84) considering q<0.2. Ten molecular surfaces were generated and averaged for each ribozyme. Scattering intensities were generated from molecular models saved during the molecular dynamics simulations (described below) and compared to experimental intensities using FOXS.(85) The model with the lowest chi-square value for RS1 and CS3 as predicted by FOXS was fit in the averaged molecular envelope using ChimeraX.(86) For RS1, the best predicted fit to the experimental data included a single model, while for CS3 the best predicted fit to the experimental data required a two-state model.

### Molecular Modeling and Simulation

Rosetta’s FARFAR2(59) method was used to generate initial molecular models for RS1 and CS3. The top 10 scoring structures were generated using secondary structure constraints derived from SHAPE and with high resolution optimization after fragment assembly using the -minimize_rna flag. For RS1, an initial set of 10 models was generated without the linker or primer region present, and the second model from this set was chosen for simulation due to the more open placement of the 5′ overhang relative to the stem-loop region (segment 1). A second set of 10 models was generated without the 5′ overhang but with the linker and primer region present. The structure of the primer, linker, and four bases upstream of the linker region were saved from the highest scoring model of this set of predictions (segment 2). Segment 2 was aligned to segment 1 based on the backbone atoms of the overlapping four 5′ bases of segment 2, which correspond to the final 3′ bases of segment 1. The coordinates of the aligned segment 2 were saved without the overlapping four 5′ bases and were joined with the coordinates of segment 1, resulting in the complete structure of RS1. For CS3, a first set of 10 models were generated that consisted of the tail and two stem-loop regions without the linker and primer, and from these model 4 was selected (segment 1). A second set of 10 models was generated that contained only the second stem-loop region, the linker, and the primer, and from these the first model was selected (segment 2). Segment 2 was aligned to segment 1 by overlapping the final 4 5′ bases of segment 1 with the first four 3′ bases of segment 2 via their backbone atoms. The coordinates of the aligned segment 2 were saved, and the coordinates of segment 1 without the second stem-loop region were saved. These two were joined to form the complete structure of CS3.

The starting structures of RS1 and CS3 used for all-atom molecular dynamics simulations were generated as described above. The VMD psfgen, solvate, and autoionize plugins were used to build both systems.(87) TIP3P water boxes were made large enough to prevent interaction with periodic images, and 50 mM NaCl was added to both systems. Both the 5’ and 3’ ends of RS1 and CS3 were capped with an -OH group. Simulations were run using NAMD 2.14(88) with the CHARMM36 force field and the TIP3P explicit model of water.(89) A cutoff of 12 Å was used for van der Waals interactions, and long-range electrostatic forces were computed using the Particle Mesh Ewald method with a grid point density of >1 Å^3^. The SHAKE algorithm was used with a timestep of 2 fs. The NpT ensemble was used at 1 atm with a hybrid Nose-Hoover Langevin piston method. The temperature was set to 300 K. Both systems were minimized for 5,000 steps, followed by 100,000 steps of equilibration in which the backbone atoms were constrained. Finally, constraints were lifted, and the system was equilibrated for 50 ns. For both RS1 and CS3, the NAMD collective variables module was used to apply weak restraints on the interactions of terminal G-C and A-U stem base pairs. Root mean square deviation (RMSD) was calculated by aligning the non-hydrogen backbone atoms to the initial frame of each simulation after constrained equilibration.

### Identification of the quasi-neutral pathway between RS1 and CS3

We employed Dijkstra’s algorithm, a common algorithm for finding the shortest paths using an adjacency matrix, to identify a path between RS1 and CS3. We used an adjacency matrix of RNA sequences obtained from all rounds of selection. The adjacency matrix was constructed by finding the Hamming distance between each pair of sequences and connecting the sequences with a distance equal to one mutation. We then ran the shortest path finding algorithm on this adjacency matrix (single mutation matrix) and looked for the path to the most similar sequence. The algorithm used for the shortest path finding was based on the breadth-first search algorithm, which is known to be efficient and accurate for finding the shortest paths in graphs.

We implemented the algorithm in Python. Our path represents the graph using a dictionary, where the keys are the nodes, and the values are the lists of adjacent nodes. The function inputs the graph, start node, and goal node and returns the shortest path between them. The function first initializes an empty list, which will track nodes that have been visited. It then uses a while loop which continues until there are no nodes left in the queue. It checks the adjacency list of nodes in the graph dictionary and iterates over each adjacent node. If the adjacent node is the goal node, the function returns the new path. Otherwise, the function moves on to the next adjacent node. If the goal node is never found, the function returns a 0, indicating that no path was found. The code begins by looping over all sequences 1 mutation from CS3 and searching for a path from RS1 to each sequence. If no complete paths are found, it starts again by looping over all sequences 2 mutations from CS3. Again, if no paths are found, it tries sequences 3 mutations from CS3 and so on until a complete path is found. Once a path is found, the code calculates the similarity between the mutant and the original CS3 using a sequence matching function. Finally, the code prints the path from RS1 to the mutant and the similarity score between the mutant and the original CS3.

We first chose the path which ended with the sequence most similar to CS3 (INT3 in Table S3). INT 3 is only 2 mutations from RS1. We then repeated the same algorithm in reverse, with CS3 as the start sequence and INT3 as the goal sequence. As expected, a compete path was not generated but the sequence closest to INT3 was revealed as INT4 which is 24 mutations from INT3 and 3 mutations from CS3. This provided us with the nearest complete path from both directions and allowed us to experimentally fill in the gaps. We designed sequence variants to complete the single-step mutational path and by experimental trial and error filled in the gaps with mutant sequences that exhibited ligase activity above our activity threshold (rate acceleration of >10% of the target sequence, CS3, i.e., 10-fold over background for AIP-ligation and 100-fold over background for PPP-ligation).

## Author Information

### Conflict of Interest

The authors declare no conflict of interest.

## Acknowledgment

We thank Drs. Filip Boskovic and Aleksander Radakovic for helpful comments on the manuscript. We also thank the MGH’s NextGen Sequencing Core for help with illumina sequencing. J.W.S. is an Investigator of the Howard Hughes Medical Institute. This work was supported in part by a grant from the Simons Foundation (290363) to J.W.S.

## Notes

### Competing Interest Statement

The authors have declared no competing interest.

