## Supplementary Information for "Evolution of the substrate specificity of an RNA ligase ribozyme from phosphorimidazole- to triphosphate-activation"

**This PDF file includes:**
Tables S1-S6
Figures S1 to S8

| | R1 | R2 | R3 | R4 | R5 | R6 |
|---|---|---|---|---|---|---|
| **Scale (pmol)** | 3000 | 1000 | 300 | 100 | 50 | 50 |
| **Reaction time (min)** | 180 | 180 | 60 | 30 | 30 | 5 |
| **[Mg$^{2+}$] (mM)** | 100 | 50 | 50 | 20 | 20 | 5 |
| **% Unique sequences in the output** | 97.6 | 94.3 | 66.1 | 7.2 | 7.2 | 6.3 |

**Table S1. Reaction parameters for each round of selection and sequencing output.** Selection was started with 3 nmol RNA to sample a large number of sequences, which was decreased to 50 pmol in the final round (R6). Reaction stringency was increased by reducing reaction times and $Mg^{2+}$ concentrations. This resulted in sequence enrichment across rounds with a marked decrease in sequence diversity in round 4 (from ~66% unique sequences to ~7% unique sequences).

| No. | Library Sequence (40 nt, 5′ → 3′) | Fractional abundance |
|---|---|---|
| | 1 5 10 15 20 25 30 35 40 | |
| 1. | ACGGGTGGGTAATCTAGTGTCCGCGGAATAGAACGAAACA | 0.540 |
| 2. | ACG**A**GTGGGTAATCTAGTGTCCGCGGAATAGAACGAAACA | 0.200 |
| 3. | ACGGGTGGGTAATCTAGTGTC**T**GCGGAATAGAACGAAACA | 0.038 |
| 4. | ACGGGTGGGTAATCT**G**GTGTCCGCGGAATAGAACGAAACA | 0.025 |
| 5. | ACGGGTGGGTAA**C**CTAGTGTCCGCGGAATAGAACGAAACA | 0.024 |
| 6. | ACGGGTG**T**GTAATCTAG**C**GTCCGCGGAATAGAACGAAACA | 0.022 |
| 7. | ACGGGTG**AA**TAATCTAG**C**GTCCGCGGAATAGAACGAAACA | 0.019 |
| 8. | ACGGGTGGGTAATCTAGTGTCCGCGGAATAGAACGA**T**ACA | 0.013 |
| 9. | ACGGGTG**A**GTAATCTAG**C**GTCCGCGGAATAGAACGAAACA | 0.011 |
| 10. | ACGGGTG**AA**TAATCTAGTGTCCGCGGAATAGAACGAAACA | 0.009 |
| 11. | ACG**A**GTGGGTAATCTAGTGTC**T**GCGGAATAGAACGAAACA | 0.008 |
| 12. | ACGGGTG**AA**TAATCTA**A**TGTCCGCGGAATAGAACGAAACA | 0.006 |
| 13. | ACG**A**GTGGGTAATCT**G**GTGTCCGCGGAATAGAACGAAACA | 0.006 |
| 14. | ACGGGTGGG**C**AATCTAGTGTCCGCGGAATAGAACGAAACA | 0.005 |
| 15. | ACG**A**GTGGGTAATCTAGTGTCCGCGGAATAGAACGA**T**ACA | 0.005 |
| 16. | ACGGGTGGGTAATCTAG**C**GTCCGCGGAATAGAACGAAACA | 0.004 |
| 17. | ACGGGTGGGTAATCTAGTGTCCGCGG**G**ATAGAACGAAACA | 0.004 |
| 18. | ACGGGTG**A**GTAATCTAGTGTCCGCGGAATAGAACGAAACA | 0.004 |
| 19. | ACGGGTG**AA**TAATCTAGTGTC**T**GCGGAATAGAACGAAACA | 0.002 |
| 20. | ACGGGTGGGTAA**C**CTAGTGTC**T**GCGGAATAGAACGAAACA | 0.002 |
| 21. | ACG**A**GTGGG**C**AATCTAGTGTCCGCGGAATAGAACGAAACA | 0.002 |
| 22. | ACG**A**GTGGGTAATCTAGTGTCCGCGG**G**ATAGAACGAAACA | 0.002 |
| 23. | ACGGGTGGGTAATCTAGTGTCCGCGGAA**G**AGAACGAAACA | 0.002 |
| 24. | ACGGGTGGGTAATCTAGT**T**TCCGCGGAATAGAACGAAACA | 0.002 |
| 25. | ACGGGTGGGTAATCTAGTGTCCGCGGA**G**TAGAACGAAACA | 0.001 |
| 26. | ACG**A**GTGGGTAA**C**CTAGTGTCCGCGGAATAGAACGAAACA | 0.001 |
| 27. | ACGGG**C**GGGTAATCTAGTGTCCGCGGAATAGAACGAAACA | 0.001 |
| 28. | ACGGGTGGGTAATCTAGTGTCCGCGGAATAGAACGAAAC**G** | 0.001 |
| 29. | ACGGGTGGGTAATCTAGTGTCCGCGGAATAGAACGAAAC**C** | 0.001 |
| 30. | ACGGGTGGGTAATCTAGTGTCCGCGGAATAGAACGA**G**ACA | 0.001 |
| 31. | ACGGGTGGGTAATCTAGTGTCCGCGGAATAGAACGAA**G**CA | 0.001 |
| 32. | ACGGGTG**T**GTAATCTAGTGTCCGCGGAATAGAACGAAACA | 0.001 |
| 33. | ACGGGTGGGTAATCTAGTGTCCGCGGAATAGA**G**CGAAACA | 0.001 |
| 34. | ACGGGTGGGTAATCTAGTG**C**CCGCGGAATAGAACGAAACA | 0.001 |
| 35. | ACGGGTG**A**GTAATCT**G**GTGTCCGCGGAATAGAACGAAACA | 0.001 |
| 36. | ACG**T**GTGGGTAATCTAGTGTCCGCGGAATAGAACGAAACA | 0.001 |
| 37. | ACGGGTGGGTAATCTAGTGTCCGCGGAATAG**G**ACGAAACA | 0.001 |
| 38. | ACGGGTGGGTAATCTAGTGTC**T**GCGGAATAGAACGA**T**ACA | 0.001 |

| No. | Library Sequence (40 nt, 5′ → 3′)<br>1 5 10 15 20 25 30 35 40 | Fractional abundance |
|---|---|---|
| 39. | ACGGGTGG**T**TAATCTAGTGTCCGCGGAATAGAACGAAACA | 0.001 |
| 40. | ACG**A**GTGGGTAATCTAG**C**GTCCGCGGAATAGAACGAAACA | 0.001 |
| 41. | ACGGGTGGGTAATC**C**AGTGTCCGCGGAATAGAACGAAACA | 0.001 |
| 42. | ACGGGTGGGTA**G**TCTAGTGTCCGCGGAATAGAACGAAACA | 0.001 |
| 43. | A**A**GGGTGGGTAATCTAGTGTCCGCGGAATAGAACGAAACA | 0.001 |
| 44. | **G**CGGGTGGGTAATCTAGTGTCCGCGGAATAGAACGAAACA | 0.001 |
| 45. | ACGGGTGGGTAATCTAGTGTCCGCG**T**AATAGAACGAAACA | 0.001 |
| 46. | ACGGGTGGGTAATCTAGTGTCCGCGGAATAGAACG**G**AACA | 0.001 |
| 47. | ACGGGTGGGTAATCTA**A**TGTCCGCGGAATAGAACGAAACA | 0.001 |
| 48. | ACGGGTGGGTAATCTAGTGTCCG**A**GGAATAGAACGAAACA | 0.001 |
| 49. | ACGGGTGGGTAATCTAGTGTCCGCGGAATAGAACG**C**AACA | 0.001 |
| 50. | ACGGGTGGGTAATCTAGTGTC**A**GCGGAATAGAACGAAACA | 0.001 |
| 51. | ACGGGTGGGTAATCTAGTGTCCGCGGAAT**G**GAACGAAACA | 0.001 |
| 52. | ACGGGTGGGTAATCT**G**GTGTCCGCGGA**G**TAGAACGAAACA | 0.001 |
| 53. | ACGGGTGGGTAATCTAGTGTC**G**GCGGAATAGAACGAAACA | 0.001 |
| 54. | ACGGGTGGGTAATCTAGTGT**T**CGCGGAATAGAACGAAACA | 0.001 |
| 55. | ACGGGTGGGTAATCT**G**GTGTCCGCGGAATAGAACGA**T**ACA | 0.001 |
| 56. | ACGGGTGGGTAATCTAGTGTC**T**GCGG**G**ATAGAACGAAACA | 0.001 |
| 57. | ACGGGTGGGTAA**C**CT**G**GTGTCCGCGGAATAGAACGAAACA | 0.001 |
| 58. | ACGGGTG**AA**TAATCTAGTGT**T**CGCGGAATAGAACGAAACA | 0.001 |
| 59. | ACG**A**GTGGGTAATCTAGT**T**TCCGCGGAATAGAACGAAACA | 0.001 |
| 60. | ACGGGTGGGTAATCTAGTGTCCGCGGAATAGAACGAAAC**T** | 0.001 |
| 61. | ACG**A**GTGGGTAATCTAGTGTCCGCGGAA**G**AGAACGAAACA | 0.001 |
| 62. | ACGGGT**A**GGTAATCTAGTGTCCGCGGAATAGAACGAAACA | 0.001 |
| 63. | ACGGGTGGGTAATCTAGTGTCC**T**CGGAATAGAACGAAACA | 0.001 |
| 64. | ACGGGTGGGTAA**C**CTAGTGTCCGCGGAATAGAACGA**T**ACA | 0.001 |
| 65. | ACGGGTG**AA**TAATCTAGTG**C**CCGCGGAATAGAACGAAACA | 0.001 |
| 66. | ACGGGTGGGTAATCTAGTGTCCGCGGAATAGAAC**T**AAACA | 0.001 |
| 67. | ACGGGTGGGTAATCTAGTGTCC**A**CGGAATAGAACGAAACA | 0.001 |
| 68. | ACGGGTGGGTAATCTAGTGTCCGCGG**T**ATAGAACGAAACA | 0.001 |

**Table S2. CS3 cluster sequences with >100 reads in round 6.** Sequences in the CS3 cluster, sorted in decreasing order of their abundance. The top 3 sequences cover ~80% of the cluster population. Nucleotide positions that differ from the peak sequence CS3 are indicated in boldface. For population dynamics across rounds, see Figure 5B.

| Name | Library Sequence (40 nt, 5′ → 3′) |
|---|---|
| | 1 5 10 15 20 25 30 35 40 |
| **RS1** | GAAUGCUGCCAACCGUGCGGGCUAAUUGGCAGACUGAGCU |
| **INT1** | GAAUGCUGCCAACCGUGCGGGCUAAUU**C**GCAGACUGAGCU |
| **INT2** | GAAUGCUGCCAACC**U**UGCGGGCUAAUU**C**GCAGACUGAGCU |
| **INT3** | GAAUGCUGCCAACC**U**UGCGGGCUAAUU**A**GCAGACUGAGCU |
| **INT4*** | **ACGA**G**UG**GC**U**AA**U**C**UA**G**U**G**UC**C**GCGGAAUAG**GAC**GA**A**A**C**A** |
| **INT5** | **ACGA**G**UG**G**GU**AA**U**C**UA**G**U**G**UC**C**GCGGAAUAG**GAC**GA**A**A**C**A** |
| **INT6** | **ACGG**G**UG**G**GU**AA**U**C**UA**G**U**G**UC**C**GCGGAAUAG**GAC**GA**A**A**C**A** |
| **CS3** | **ACGG**G**UG**G**GU**AA**U**C**UA**G**U**G**UC**C**GCGGAAUAGA**AC**GA**A**A**C**A** |

**Table S3. Intermediate sequences between RS1 and CS3, identified by mining high-throughput sequencing data from rounds 1-6.** INT4 (indicated by an asterisk) resembles CS3 (3 mutations from it) and is 25 mutations from RS1. The mutational space between INT3 and INT4 was filled experimentally to generate Figure 6 and Table S4.

| Name | Library Sequence (40 nt, 5′ → 3′) | $k_{obs}$ ($h^{-1}$) AIP | $k_{obs}$ ($h^{-1}$) PPP |
|---|---|---|---|
| | 1 5 10 15 20 25 30 35 40 | | |
| RS1 | GAAUGCUGCCAACCGUGCGGGCUAAUUGGCAGACUGAGCU | 6.38 ± 0.60 | - |
| Int1 | GAAUGCUGCCAACC**U**UGCGGGCUAAUUGGCAGACUGAGCU | 1.78 ± 0.24 | - |
| Int2 | GAAUGCUGCCAACC**U**UGCGGGCUAAUU**A**GCAGACUGAGCU | 2.78 ± 0.36 | - |
| Int3 | GAA**A**GCUGCCAACC**U**UGCGGGCUAAUU**A**GCAGACUGAGCU | 1.59 ± 0.23 | - |
| Int4 | GAA**A**GCUGC**U**AACC**U**UGCGGGCUAAUU**A**GCAGACUGAGCU | 0.30 ± 0.08 | - |
| Int5 | GAA**A**GCUGC**U**AACC**UA**GCGGGCUAAUU**A**GCAGACUGAGCU | 0.19 ± 0.04 | - |
| Int6 | GAA**A**GCUGC**U**AACC**UA**GCGG**C**CUAAUU**A**GCAGACUGAGCU | 0.20 ± 0.05 | - |
| Int7 | GAA**A**GCUGC**U**AACC**UA**GCGG**C**CUAAUU**A**GCAGACU**A**AGCU | 0.103 ± 0.01 | - |
| Int8 | GAA**A**GCUGC**U**AACC**UA**GCGG**C**CUAAUU**A**G**A**AGACU**A**AGCU | 0.134 ± 0.12 | - |
| Int9 | GAA**A**GCUGC**U**AA**U**C**UA**GCGG**C**CUAAUU**A**G**A**AGACU**A**AGCU | 0.20 ± 0.03 | - |
| Int10 | GAA**A**GCUGC**U**AA**U**C**UA**GCGG**C**C**G**AAUU**A**G**A**AGACU**A**AGCU | 0.30 ± 0.06 | - |
| Int11 | GAA**A**GCUGC**U**AA**U**C**UA**GCGG**C**C**G**AAUU**A**G**A**AGACU**A**AGC**A** | 0.12 ± 0.04 | - |
| Int12 | GAA**A**GCUGC**U**AA**U**C**UA**GCGG**C**C**G**AAUU**A**G**A**AGACU**A**A**A**C**A** | 0.38 ± 0.05 | - |
| Int13 | GAA**A**GCUGC**U**AA**U**C**UA**GCGG**C**C**G**AAUU**A**G**A**AGAC**GA**A**A**C**A** | 0.61 ± 0.08 | - |
| Int14 | GAA**A**GCUGC**U**AA**U**C**UA**GCGG**C**C**G**AAUU**A**G**AG**GAC**GA**A**A**C**A** | 0.66 ± 0.19 | - |
| Int15 | GAA**A**GCUGC**U**AA**U**C**UA**GCGG**C**C**G**AAUU**AUAG**GAC**GA**A**A**C**A** | 0.26 ± 0.04 | - |
| Int16 | GAA**A**GCUGC**U**AA**U**C**UA**GCGG**C**C**G**AAU**AAUAG**GAC**GA**A**A**C**A** | 0.11 ± 0.05 | - |
| Int17 | GAA**A**GC**G**GC**U**AA**U**C**UA**GCGG**C**C**G**AAU**AAUAG**GAC**GA**A**A**C**A** | 0.09 ± 0.05 | - |
| Int18 | GAA**A**GC**G**GC**U**AA**U**C**UA**GCGG**C**C**G**AA**GAAUAG**GAC**GA**A**A**C**A** | 0.12 ± 0.04 | - |
| Int19 | GAA**A**GC**G**GC**U**AA**U**C**UA**GCGG**C**C**G**A**GGAAUAG**GAC**GA**A**A**C**A** | 0.09 ± 0.05 | - |
| Int20 | GAA**A**GC**G**GC**U**AA**U**C**UA**GCGG**C**C**GCGGAAUAG**GAC**GA**A**A**C**A** | 1.00 ± 0.05 | - |
| Int21 | GAA**A**GC**G**GC**U**AA**U**C**UA**GCG**UC**C**GCGGAAUAG**GAC**GA**A**A**C**A** | 0.25 ± 0.04 | - |
| Int22 | GAA**A**GC**G**GC**U**AA**U**C**UA**G**U**G**UC**C**GCGGAAUAG**GAC**GA**A**A**C**A** | 0.46 ± 0.01 | - |
| Int23 | GAA**A**G**UG**GC**U**AA**U**C**UA**G**U**G**UC**C**GCGGAAUAG**GAC**GA**A**A**C**A** | 0.21 ± 0.02 | - |
| Int24 | GA**GA**G**UG**GC**U**AA**U**C**UA**G**U**G**UC**C**GCGGAAUAG**GAC**GA**A**A**C**A** | 0.14 ± 0.05 | - |
| Int25 | G**CGA**G**UG**GC**U**AA**U**C**UA**G**U**G**UC**C**GCGGAAUAG**GAC**GA**A**A**C**A** | 0.19 ± 0.05 | - |
| Int26 | **ACGA**G**UG**GC**U**AA**U**C**UA**G**U**G**UC**C**GCGGAAUAG**GAC**GA**A**A**C**A** | 1.03 ± 0.15 | 0.009 ± 0.002 |
| Int27 | **ACGA**G**UG**GC**U**AA**U**C**UA**G**U**G**UC**C**GCGGAAUAGA**AC**GA**A**A**C**A** | 0.86 ± 0.13 | 0.004 ± 0.001 |
| Int28 | **ACGA**G**UG**G**GU**AA**U**C**UA**G**U**G**UC**C**GCGGAAUAGA**AC**GA**A**A**C**A** | 0.45 ± 0.01 | 0.005 ± 0.001 |
| CS3 | **ACGG**G**UG**G**GU**AA**U**C**UA**G**U**G**UC**C**GCGGAAUAGA**AC**GA**A**A**C**A** | 1.06 ± 0.01 | 0.012 ± 0.003 |
| | 1 5 10 15 20 25 30 35 40 | | |

**Table S4. Mutational pathway connecting RS1 to CS3.** Point mutations to RS1 are highlighted in bold. PPP-Ligase activity emerges in Sequence 26, which is accompanied by a marked increase in AIP-ligation. The two-dimensional fitness landscape for ligase function containing these sequences is shown in Figure 6.

| SAXS parameters | RS1 | CS3 |
|---|---|---|
| $R_g$ (Å) [from P(r)] | 36.07 ± 1.81 | 37.93 ± 0.94 |
| $R_g$ (Å) [from Guinier] | 30.33 ± 1.34 | 31.38 ± 3 |
| $D_{max}$ (Å) | 149.0 | 123.0 |
| Molecular Weight (kDa) | 32.8 | 36.4 |

**Table S5. SAXS Parameters for RS1 and CS3.**

| No. | Oligo Name | Sequence (5'→3') | Type | Source |
|---|---|---|---|---|
| 1.1 | RS1 | GACUCACUGACACAGAUCCACUCACGGACAGCG GAAUGCUGCCAACCGUGCGGGCUAAUUGGCAGA CUGAGCUCGCUGUCC*UUUUUU*GGCUAAG**G** | RNA | IVT |
| 1.2 | RS1_doped21_DNA (Mutagenesis at 21% at each nucleotide position: 79% WT nucleotide and 7% of the other three) Note: The sequence is written according to IDT specifications | TAATACGACTCACTATAGACTCACTGACACAGA TCCACTCACGGACAGCG(N1:07077907)(N2:79070707)(N2)(N3:07070779)(N1)(N4:07790707)(N3)(N1)(N4)(N4)(N2)(N2)(N4)(N4)(N1)(N3)(N1)(N4)(N1)(N1)(N1)(N4)(N3)(N2)(N2)(N3)(N3)(N1)(N1)(N4)(N2)(N1)(N2)(N4)(N3)(N1)(N2)(N1)(N4)(N3)CGCTGTCC*TTTTTT*GGCT AAG**G** | DNA | IDT |
| 2.1 | Template | GCGGUGGUCCUUAGCC | RNA | IDT |
| 2.2 | Template+1 | UGCGGUGGUCCUUAGCC | RNA | IDT |
| 2.3 | Template+2 | AUGCGGUGGUCCUUAGCC | RNA | IDT |
| 2.4 | Template+3 | AAUGCGGUGGUCCUUAGCC | RNA | IDT |
| 2.5 | Template+4 | GAAUGCGGUGGUCCUUAGCC | RNA | IDT |
| 2.6 | Template+5 | GGAAUGCGGUGGUCCUUAGCC | RNA | IDT |
| 2.7 | Template+6 | CGGAAUGCGGUGGUCCUUAGCC | RNA | IDT |
| 2.8 | Template+7 | GCGGAAUGCGGUGGUCCUUAGCC | RNA | IDT |
| 2.9 | Template+8 | UGCGGAAUGCGGUGGUCCUUAGCC | RNA | IDT |
| 2.10 | Template_U8A | GCGGUGGACCUUAGCC | RNA | IDT |
| 2.11 | Template_C9G | GCGGUGGUGCUUAGCC | RNA | IDT |
| 3.1 | PPP-LigB | (5'-triphosphate) - ACCACCGCAUUCCGCA - (3BioTEG) | RNA | Chemgenes |
| 3.2 | PPP-Lig | (5'-triphosphate) - ACCACCGCAUUCCGCA | RNA | Chemgenes |
| 3.3 | AIP-Lig | (5'-phosphoro-2-aminoimidazole) - ACCACCGCAUUCCGCA | RNA | Activation of P-Lig |
| 3.4 | P-LigB | (5'-monophosphate) - ACCACCGCAUUCCGCA - (3BioTEG) | RNA | IDT |
| 3.5 | P-Lig | (5'-monophosphate) - ACCACCGCAUUCCGCA | RNA | IDT |
| 3.6 | HO-Lig | (5'-hydroxyl) - ACCACCGCAUUCCGCA | RNA | IDT |
| 3.7 | MeIP-Lig | (5'-phosphoro-2-methylimidazole) - ACCACCGCAUUCCGCA | RNA | Activation of P-Lig |
| 4 | RT primer | GTGCGGAATGCGGTGGTCCTT | DNA | IDT |
| 5.1 | Quench oligo 1 | GTGCGGAATGCGGTGGT | DNA | IDT |
| 5.2 | Quench oligo 2 | GCCTTAGCCAAAAAAGGACAGCG | DNA | IDT |
| 6.1 | PCR_Forward_primer | TAATACGACTCACTATAGACTCACTGACAC | DNA | IDT |
| 6.2 | PCR_Reverse_primer | mCmCTTAGCCAAAAAAGGACAGCG | DNA | IDT |
| 6.3 | SeqPCR_primer1 | GTTCAGAGTTCTACAGTCCGACGATCCGGTAGG TCCCTTAGCCAAAAAAGGACAGCG | DNA | IDT |
| 6.4 | SeqPCR_primer2 | AGACGTGTGCTCTTCCGATCTGACTCACTGACA CAGATCCACTCAC | DNA | IDT |
| 6.5 | SeqPCR_primer3 | GTTCAGAGTTCTACAGTCCGACGATC | DNA | IDT |
| 6.6 | SeqPCR_primer4 | AGACGTGTGCTCTTCCGATCT | DNA | IDT |
| 7.1 | CS1 | GACUCACUGACACAGAUCCACUCACGGACAGCG GACAGCCGAGAAAUGAGUGGCCUAAAUGGGAGA AUGAGCUCGCUGUCC*UUUUUU*GGCUAAG**G** | RNA | IVT |

| | | | | |
|---|---|---|---|---|
| 7.2 | CS2 | GACUCACUGACACAGAUCCACUCACGGACAGCG GACUGCGCGUAUGAGUGGCGGCUAAAGAGGAGA AUGAGCGCGCUGUCC*UUUUUU*GGCUAAG**G** | RNA | IVT |
| 7.3 | CS3 | GACUCACUGACACAGAUCCACUCACGGACAGCG ACGGGUGGGUAAUCUAGUGUCCGCGGAAUAGAA CGAAACACGCUGUCC*UUUUUU*GGCUAAG**G** | RNA | IVT |
| 7.4 | CS3_5't | GGACAGCGACGGGUGGGUAAUCUAGUGUCCGCG GAAUAGAACGAAACACGCUGUCC*UUUUUU*GGCU AAG**G** | RNA | IVT |
| 7.5 | CS3_3't | GACUCACUGACACAGAUCCACUCACGGACAGCG ACGGGUGGGUAAUCUAGUGUCCGCGGAAUAGAA CGAAACACGCUGUCC | RNA | IVT |
| 7.6 | CS4 | GACUCACUGACACAGAUCCACUCACGGACAGCG GGAUGGUGCGAACUGAGUGGGCUAAUUAGGAGA AUGAGCGCGCUGUCC*UUUUUU*GGCUAAG**G** | RNA | IVT |
| 7.7 | CS5 | GACUCACUGACACAGAUCCACUCACGGACAGCG GGAGGGUGACAUCGUUGAGAGAGAAUGGGGAUA UUGAACUCGCUGUCC*UUUUUU*GGCUAAG**G** | RNA | IVT |
| 8.1 | PPP-CS3_pc1 | (5'-triphosphate) - GACUCACUGACACAGAUCCACUCAC | RNA | IVT |
| 8.2 | P-CS3_pc1 | (5'-monophosphate) - GACUCACUGACACAGAUCCACUCAC | RNA | IDT |
| 8.3 | HO-CS3_pc1 | (5'-hydroxyl) - GACUCACUGACACAGAUCCACUCAC | RNA | IDT |
| 8.4 | pCS3_pc2 | (5'-monophosphate) - GGACAGCGACGGGUGGGUAAUCUAGUGUCCGCG GAAUAGA | RNA | IDT |
| 8.5 | pCS3_pc3 | (5'-monophosphate) - ACGAAACACGCUGUCCUUUUUUGGCUAAGG | RNA | IDT |
| 8.6 | CS3_splint1 | ACCCACCCGTCGCTGTCCGTGAGTGGATCTGTG TCAG | DNA | IDT |
| 8.7 | CS3_splint2 | CCAAAAAAGGACAGCGTGTTTCGTTCTATTCCG CGGACAC | DNA | IDT |
| 9.1 | CS3_SHAPE | GGCCUUCGGGCCAAGACUCACUGACACAGAUCC ACUCACGGACAGCGACGGGUGGGUAAUCUAGUG UCCGCGGAAUAGAACGAAACACGCUGUCCUUUU UUGGCUAAG**G**UCGAUCCGGUUCGCCGGAUCCAA AUCGGGCUUCGGUCCGGUUC | RNA | IVT |
| 9.2 | SHAPE_RT_primer | (5'-FAM)-GAACCGGACCGAAGCCCG | | |
| 10.1 | Int1 | GACUCACUGACACAGAUCCACUCACGGACAGCG GAAUGCUGCCAACCUUGCGGGCUAAUUGGCAGA CUGAGCUCGCUGUCC*UUUUUU*GGCUAAG**G** | RNA | IVT |
| 10.2 | Int2 | GACUCACUGACACAGAUCCACUCACGGACAGCG GAAUGCUGCCAACCUUGCGGGCUAAUUAGCAGA CUGAGCUCGCUGUCC*UUUUUU*GGCUAAG**G** | RNA | IVT |
| 10.3 | Int3 | GACUCACUGACACAGAUCCACUCACGGACAGCG GAAAGCUGCCAACCUUGCGGGCUAAUUAGCAGA CUGAGCUCGCUGUCC*UUUUUU*GGCUAAG**G** | RNA | IVT |
| 10.4 | Int4 | GACUCACUGACACAGAUCCACUCACGGACAGCG GAAAGCUGCUAACCUUGCGGGCUAAUUAGCAGA CUGAGCUCGCUGUCC*UUUUUU*GGCUAAG**G** | RNA | IVT |
| 10.5 | Int5 | GACUCACUGACACAGAUCCACUCACGGACAGCG GAAAGCUGCUAACCUAGCGGGCUAAUUAGCAGA CUGAGCUCGCUGUCC*UUUUUU*GGCUAAG**G** | RNA | IVT |

| | | | | |
|---|---|---|---|---|
| 10.6 | Int6 | GACUCACUGACACAGAUCCACUCACGGACAGCG<br>GAAAGCUGCUAACCUAGCGGCCUAAUUAGCAGA<br>CUGAGCUCGCUGUCC*UUUUUU*GGCUAAG**G** | RNA | IVT |
| 10.7 | Int7 | GACUCACUGACACAGAUCCACUCACGGACAGCG<br>GAAAGCUGCUAACCUAGCGGCCUAAUUAGCAGA<br>CUAAGCUCGCUGUCC*UUUUUU*GGCUAAG**G** | RNA | IVT |
| 10.8 | Int8 | GACUCACUGACACAGAUCCACUCACGGACAGCG<br>GAAAGCUGCUAACCUAGCGGCCUAAUUAGAAGA<br>CUAAGCUCGCUGUCC*UUUUUU*GGCUAAG**G** | RNA | IVT |
| 10.9 | Int9 | GACUCACUGACACAGAUCCACUCACGGACAGCG<br>GAAAGCUGCUAAUCUAGCGGCCUAAUUAGAAGA<br>CUAAGCUCGCUGUCC*UUUUUU*GGCUAAG**G** | RNA | IVT |
| 10.10 | Int10 | GACUCACUGACACAGAUCCACUCACGGACAGCG<br>GAAAGCUGCUAAUCUAGCGGCCGAAUUAGAAGA<br>CUAAGCUCGCUGUCC*UUUUUU*GGCUAAG**G** | RNA | IVT |
| 10.11 | Int11 | GACUCACUGACACAGAUCCACUCACGGACAGCG<br>GAAAGCUGCUAAUCUAGCGGCCGAAUUAGAAGA<br>CUAAGCACGCUGUCC*UUUUUU*GGCUAAG**G** | RNA | IVT |
| 10.12 | Int12 | GACUCACUGACACAGAUCCACUCACGGACAGCG<br>GAAAGCUGCUAAUCUAGCGGCCGAAUUAGAAGA<br>CUAAACACGCUGUCC*UUUUUU*GGCUAAG**G** | RNA | IVT |
| 10.13 | Int13 | GACUCACUGACACAGAUCCACUCACGGACAGCG<br>GAAAGCUGCUAAUCUAGCGGCCGAAUUAGAAGA<br>CGAAACACGCUGUCC*UUUUUU*GGCUAAG**G** | RNA | IVT |
| 10.14 | Int14 | GACUCACUGACACAGAUCCACUCACGGACAGCG<br>GAAAGCUGCUAAUCUAGCGGCCGAAUUAGAGGA<br>CGAAACACGCUGUCC*UUUUUU*GGCUAAG**G** | RNA | IVT |
| 10.15 | Int15 | GACUCACUGACACAGAUCCACUCACGGACAGCG<br>GAAAGCUGCUAAUCUAGCGGCCGAAUUAUAGGA<br>CGAAACACGCUGUCC*UUUUUU*GGCUAAG**G** | RNA | IVT |
| 10.16 | Int16 | GACUCACUGACACAGAUCCACUCACGGACAGCG<br>GAAAGCUGCUAAUCUAGCGGCCGAAUAAUAGGA<br>CGAAACACGCUGUCC*UUUUUU*GGCUAAG**G** | RNA | IVT |
| 10.17 | Int17 | GACUCACUGACACAGAUCCACUCACGGACAGCG<br>GAAAGCGGCUAAUCUAGCGGCCGAAUAAUAGGA<br>CGAAACACGCUGUCC*UUUUUU*GGCUAAG**G** | RNA | IVT |
| 10.18 | Int18 | GACUCACUGACACAGAUCCACUCACGGACAGCG<br>GAAAGCGGCUAAUCUAGCGGCCGAAGAAUAGGA<br>CGAAACACGCUGUCC*UUUUUU*GGCUAAG**G** | RNA | IVT |
| 10.19 | Int19 | GACUCACUGACACAGAUCCACUCACGGACAGCG<br>GAAAGCGGCUAAUCUAGCGGCCGAGGAAUAGGA<br>CGAAACACGCUGUCC*UUUUUU*GGCUAAG**G** | RNA | IVT |
| 10.20 | Int20 | GACUCACUGACACAGAUCCACUCACGGACAGCG<br>GAAAGCGGCUAAUCUAGCGGCCGCGGAAUAGGA<br>CGAAACACGCUGUCC*UUUUUU*GGCUAAG**G** | RNA | IVT |
| 10.21 | Int21 | GACUCACUGACACAGAUCCACUCACGGACAGCG<br>GAAAGCGGCUAAUCUAGCGUCCGCGGAAUAGGA<br>CGAAACACGCUGUCC*UUUUUU*GGCUAAG**G** | RNA | IVT |
| 10.22 | Int22 | GACUCACUGACACAGAUCCACUCACGGACAGCG<br>GAAAGCGGCUAAUCUAGUGUCCGCGGAAUAGGA<br>CGAAACACGCUGUCC*UUUUUU*GGCUAAG**G** | RNA | IVT |

| | | | | |
|---|---|---|---|---|
| 10.23 | Int23 | GACUCACUGACACAGAUCCACUCACGGACAGCG GAAAGUGGCUAAUCUAGUGUCCGCGGAAUAGGA CGAAACACGCUGUCC*UUUUUU*GGCUAAG**G** | RNA | IVT |
| 10.24 | Int24 | GACUCACUGACACAGAUCCACUCACGGACAGCG GAGAGUGGCUAAUCUAGUGUCCGCGGAAUAGGA CGAAACACGCUGUCC*UUUUUU*GGCUAAG**G** | RNA | IVT |
| 10.25 | Int25 | GACUCACUGACACAGAUCCACUCACGGACAGCG GCGAGUGGCUAAUCUAGUGUCCGCGGAAUAGGA CGAAACACGCUGUCC*UUUUUU*GGCUAAG**G** | RNA | IVT |
| 10.26 | Int26 | GACUCACUGACACAGAUCCACUCACGGACAGCG ACGAGUGGCUAAUCUAGUGUCCGCGGAAUAGGA CGAAACACGCUGUCC*UUUUUU*GGCUAAG**G** | RNA | IVT |
| 10.27 | Int27 | GACUCACUGACACAGAUCCACUCACGGACAGCG ACGAGUGGCUAAUCUAGUGUCCGCGGAAUAGAA CGAAACACGCUGUCC*UUUUUU*GGCUAAG**G** | RNA | IVT |
| 10.28 | Int28 | GACUCACUGACACAGAUCCACUCACGGACAGCG ACGAGUGGGUAAUCUAGUGUCCGCGGAAUAGAA CGAAACACGCUGUCC*UUUUUU*GGCUAAG**G** | RNA | IVT |

**Table S6. Oligonucleotides used in this work.** In the ribozyme sequences, randomized nucleotides are highlighted in blue, the T7 promoter sequence is shown in purple, the $U_6$ linker is shown in italics, the primer sequence is underlined, and the 3′ terminal nucleotide possessing the nucleophile for the ligation reaction is shown in boldface. 5′ and 3′ SHAPE cassettes are highlighted in green. Oligonucleotides were either purchased from Integrated DNA Technologies (IDT) or generated enzymatically by *in vitro* transcription (IVT) of dsDNA templates. AIP-Lig and MeIP-Lig were generated by incubating P-Lig with EDC and 2-aminoimidazole (2AI) and 2-methylamidazole (2MeI), respectively (See 'RNA preparation and substrate activation' in Materials and Methods).

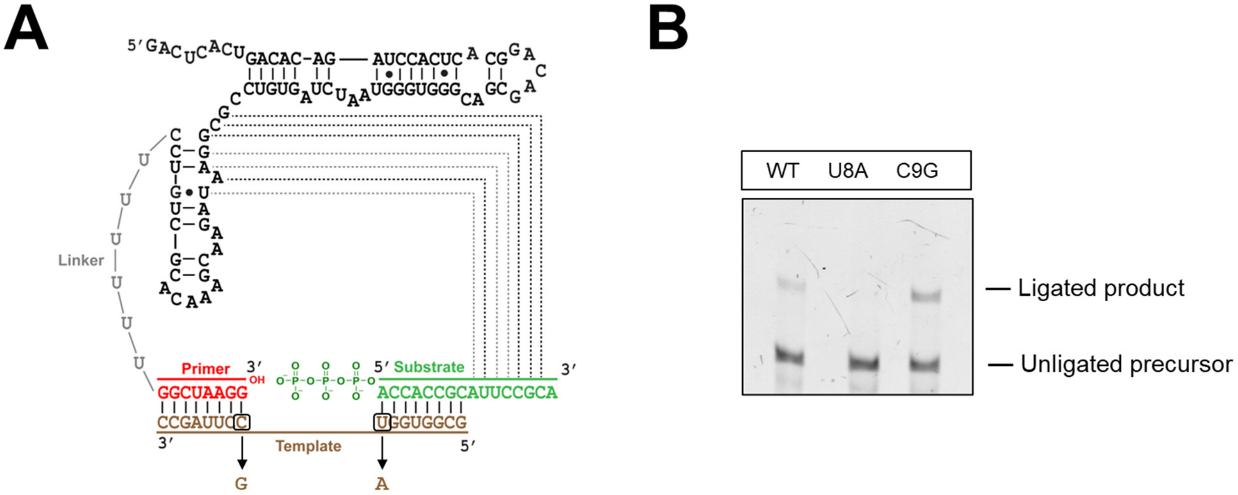


**Figure S1. Effect of disrupting the ligation junction. A.** CS3 in complex with the template and substrate, showing mutations in the template (U8A and C9G) that disrupt the ligation junction at the substrate and primer end, respectively. **B.** The U8A mutation, which unpairs the terminal base pair between the substrate and the template, abrogates ligation. The C9G mutation, which unpairs the terminal base pair between the primer and the template, preserves ligation. Ligation reactions contained 1 μM ribozyme, 1.2 μM RNA template, and 2 μM PPP-Lig in 100 mM Tris-HCl (pH 8.0), 300 mM NaCl, and 100 mM $MgCl_2$. Template sequences used in these experiments can be found in Table S6.

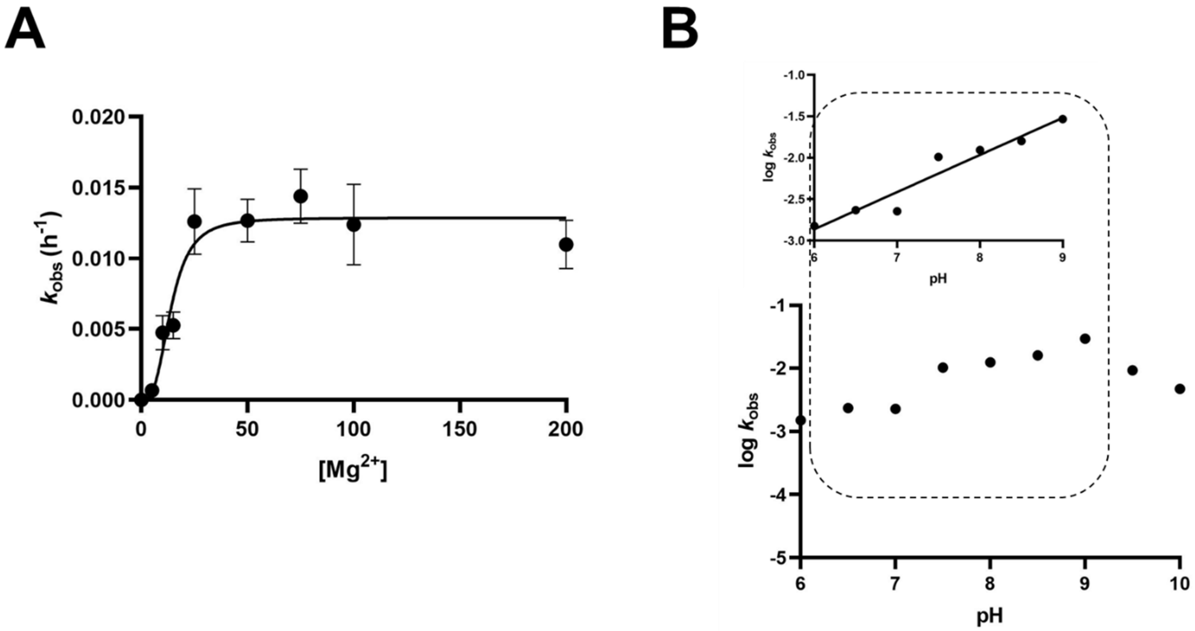


**Figure S2. The effect of $Mg^{2+}$ concentration and pH on the rates of CS3-catalyzed ligation of PPP-Lig. A.** The $k_{obs}$ value for PPP-ligation increase steeply till [$Mg^{2+}$] reaches 25 mM and plateaus at higher concentrations. The data is plotted to the Hill Equation and yields a $[Mg^{2+}]_{1/2}$ = 13.9 ± 1.79 mM and is consistent with the binding of ~3 $Mg^{2+}$ ions. Ligation reactions contained 1 μM ribozyme, 1.2 μM RNA template, and 2 μM PPP-Lig in 100 mM Tris-HCl (pH 8.0), 300 mM NaCl, and the indicated amounts of $MgCl_2$. **B.** Ligation rates increase from pH 6 to pH 9 and then fall. The inset shows that the increase in ligation rate is log-linear in the range pH 6-9 with a slope of 0.45 ± 0.05. Ligation conditions were identical as in (A), except that the reactions contained 100 mM $MgCl_2$ and were performed at the indicated pH values.

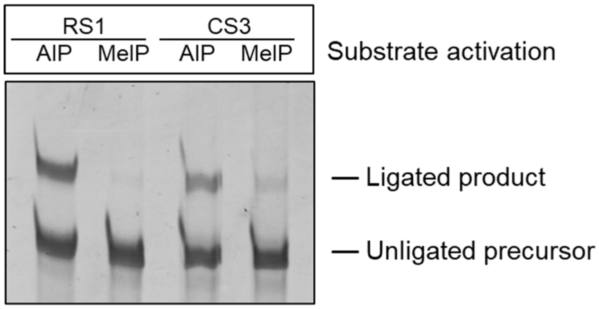


**Figure S3. The effect of substrate activation on ligation by RS1 and CS3**. RS1 is specific to 2-aminoimidazole-activated substrates; however, CS3 can ligate substrates activated with both 2-aminoimidazole (AIP-Lig) and 2-methylimidazole (MeIP-Lig). Ligation reactions contained 1 μM ribozyme, 1.2 μM RNA template, and 2 μM substrate in 100 mM Tris-HCl (pH 8.0), 300 mM NaCl, and 10 mM $MgCl_2$.

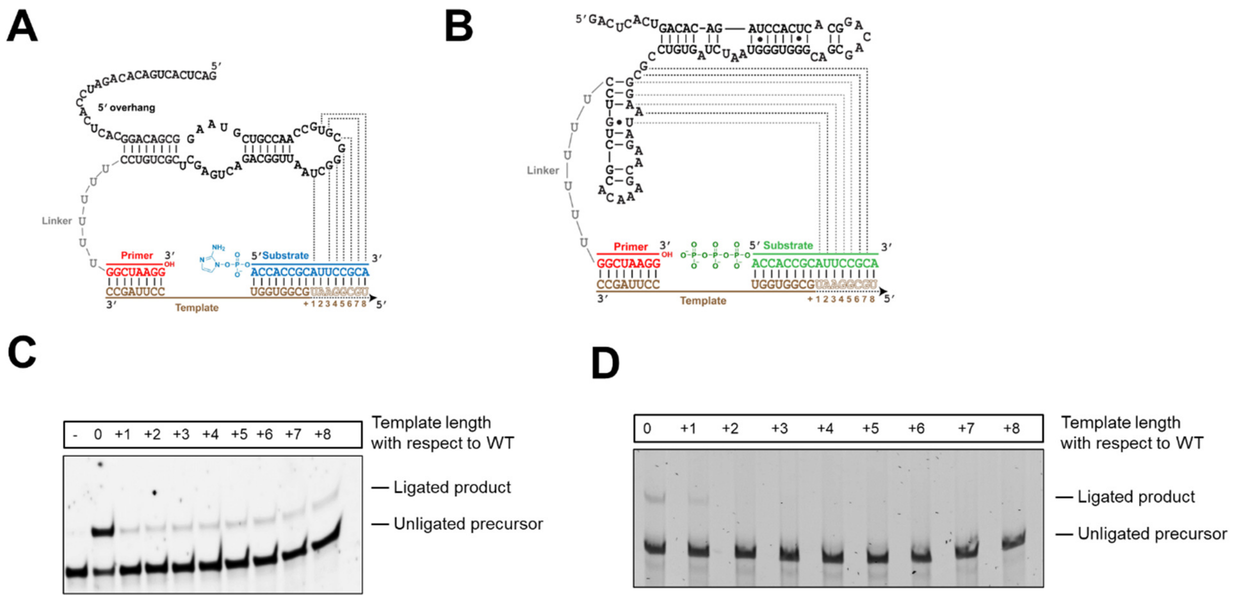


**Figure S4. Progressive sequestering of the 3′ substrate overhang results in a decrease in ligation by RS1 and CS3.** Extending the length of the template (brown) at its 5′ end sequesters the 3' overhang of **A.** AIP-Lig and **B.** PPP-Lig. **C.** Significant reduction in ligation is observed when the template is extended by 1 nt in the case of AIP-ligation. **D.** Extending the template by 1 nt reduces ligation for PPP-ligation, and further extension eliminates ligation. Ligation reactions contained 1 μM ribozyme, 1.2 μM RNA template, and 2 μM AI-Lig or PPP-Lig in 100 mM Tris-HCl (pH 8.0), 300 mM NaCl, and 10 mM (for AIP-ligation) or 100 mM (for PPP-ligation) $MgCl_2$. Template sequences used in these experiments can be found in Table S6.

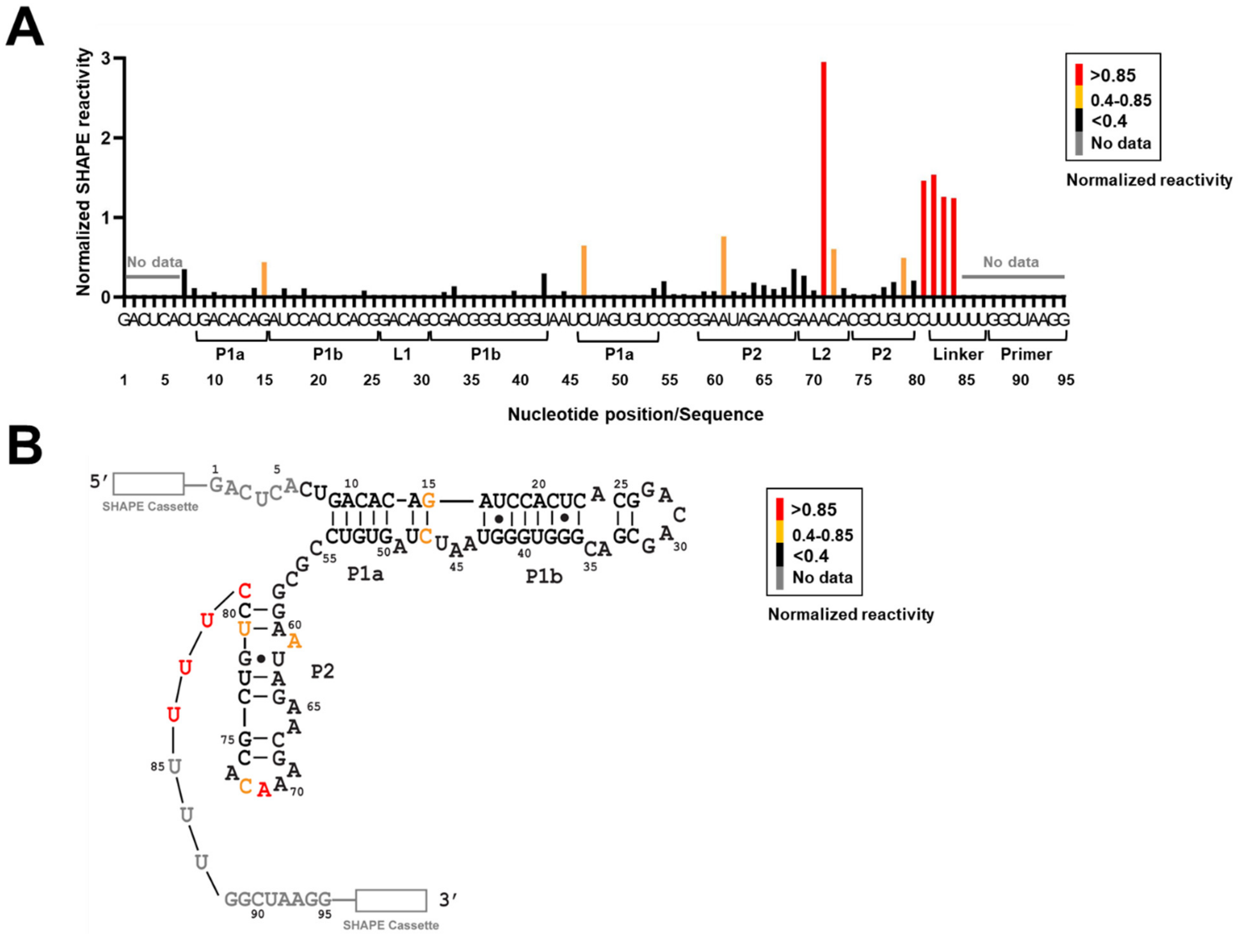


**Figure S5. SHAPE analysis of CS3.** Products of primer extension reactions with FAM-labeled primers after modification with 1M7 were separated on a 10% denaturing PAGE. **A.** Normalized SHAPE reactivities were plotted for each nucleotide position. High reactivity suggests flexibility within the RNA structure, while low reactivity suggests base-paired stems or interactions with distal nucleotides in its tertiary fold. **B.** Secondary structure of CS3 determined by the RNAstructure program using reactivity constraints obtained from SHAPE experiments. 5′ and 3′ SHAPE cassettes are denoted by white rectangles. Nucleotides for which no data was obtained are shown in gray. Nucleotides are colored in red, orange, and black according to their normalized SHAPE reactivities, as shown in the reactivity legend. Sequences used in SHAPE experiments can be found in Table S6.

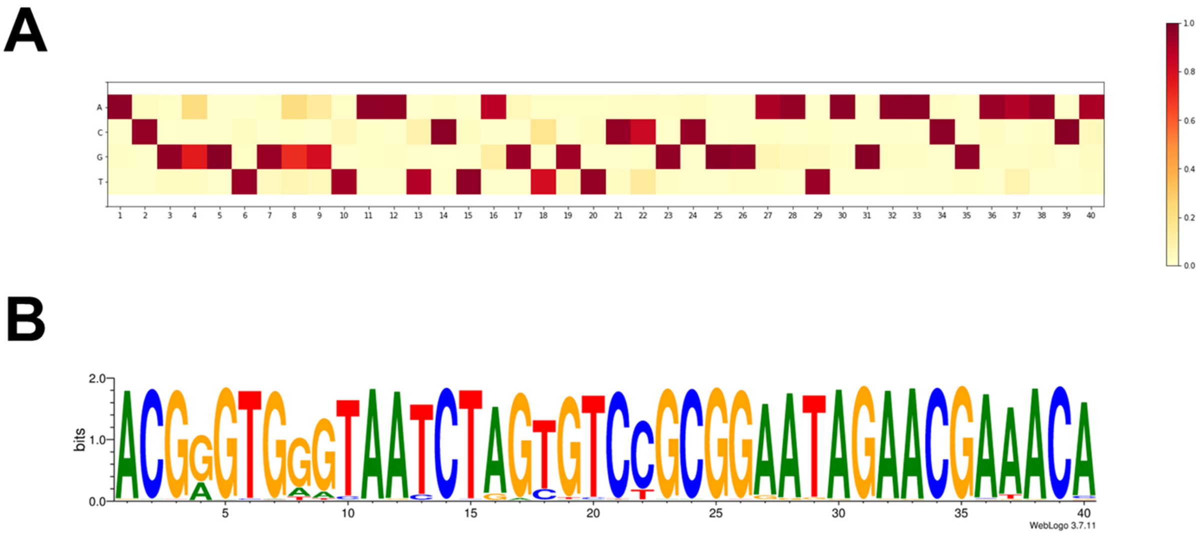


**Figure S6. High degree of nucleotide conservation in the 40 nt variable region of CS3. A.** Heat map showing the fractional abundances of each of the four nucleotides for each nucleotide position. **B.** A sequence logo shows the consensus sequence, depicting the relative abundances of each nucleotide for each position. The sequence logo was generated by the WebLogo online server (https://weblogo.berkeley.edu/logo.cgi).

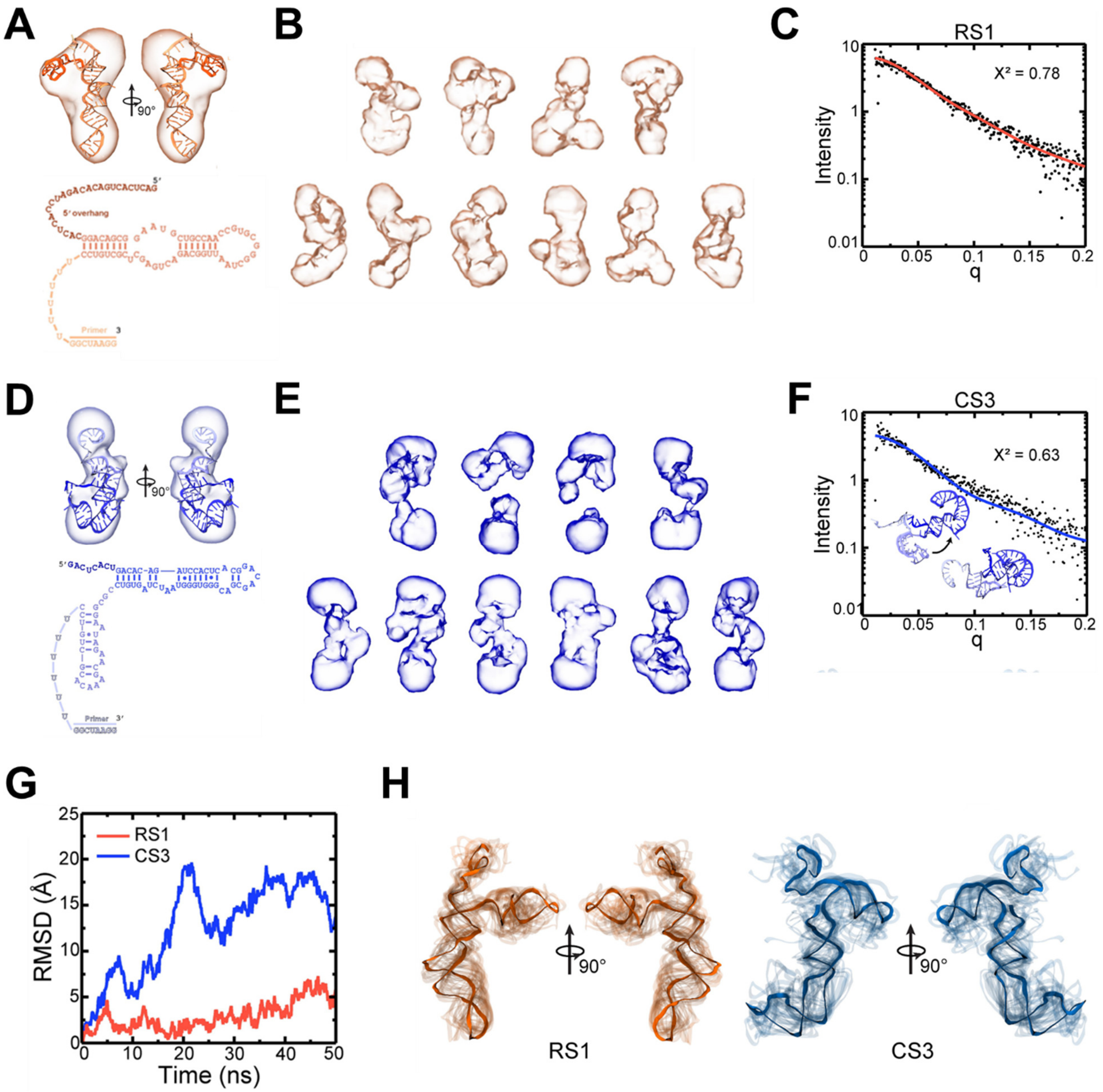


**Figure S7. Structural modeling of RS1 and CS3. A.** The computed molecular model of RS1 is shown fit to the calculated molecular envelope, with the color gradient of the molecular model corresponding to the secondary structure diagram. **B.** The ten surfaces shown were calculated by DENSS and averaged to give the final envelope in (A). **C.** Computed intensity profile (orange line) to the experimental SAXS profile (black dots) of the molecular model exhibiting the lowest chi-square value. **D.** The computed molecular model of CS3 is shown fit to the calculated molecular envelope, with the color gradient of the molecular model corresponding to the secondary structure diagram. **E.** The ten surfaces shown were calculated by DENNS and averaged to give the final envelope in (D). **F.** Computed intensity profile (blue line) to the experimental SAXS profile (black dots) of the molecular models exhibiting the best fit. The two models included in this calculation are shown in the inset. **G.** RMSD values shown for RS1 and

CS3 over the 50 ns equilibrations. **H.** Structures of RS1 and CS3 were aligned by their non-hydrogen backbone atoms and are shown every 2 ns in transparent backbone representation. The final frame of the simulation is shown in opaque backbone representation. The RMSD and molecular figures indicate a more dynamic structure for CS3, which agrees with the two-state fit to the SAXS data for CS3.

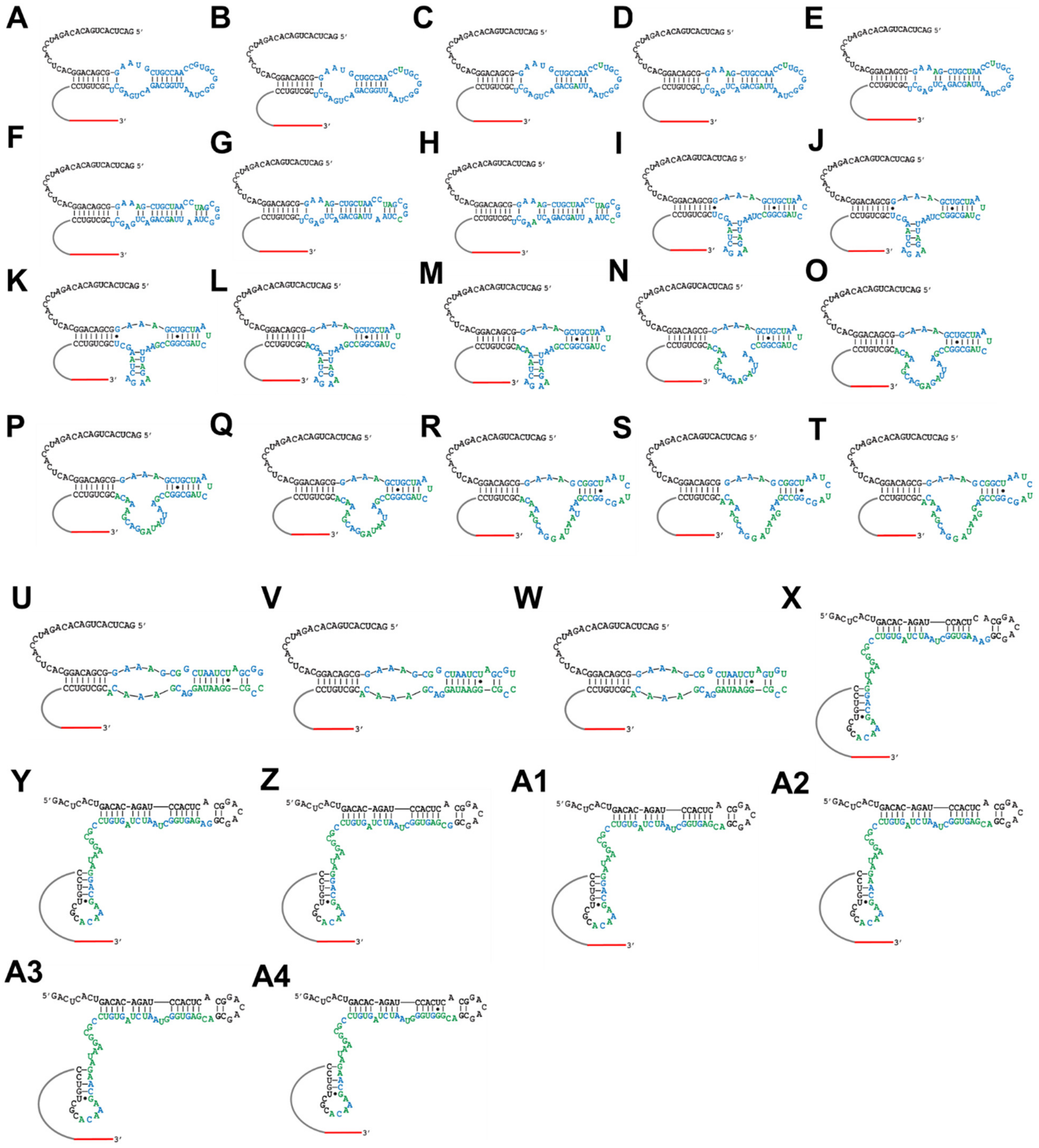


**Figure S8. Computationally predicted secondary structures of active ligase intermediates Int1-Int28 that connect RS1 and CS3 via a single-step mutational pathway.** AIP-Ligase, RS1 (A) and PPP-Ligase, CS3 (A4) are connected via mutational intermediates Int1-Int28 (B-A3) that collectively constitute a quasi-neutral pathway (see Figure 6, Table S4), enabling smooth interconversion between the two. The $U_6$ linker and 3′ primer sequences are depicted as gray and red lines. Invariant nucleotides are shown in black. The partially randomized region of RS1 is shown in blue in (A). Mutations to RS1 are shown in green. The 28 intermediate sequences can be approximately classified into 7 structure-types. A-E (RS1-like), F-H, I-M, N-Q, R-T, U-W, and X-A4 (CS3-like). Structural changes are gradual before Int23 (X), with a notable absence of base-pairing between nucleotides from the constant (shown in black) and variable (shown in blue and green) regions. The CS3-like secondary structure emerges suddenly as a result of a single mutation to Int22 (W) and is significantly different from it. In contrast to its precursors, the CS3-like structure of Int23-Int28 (X-A3) is created by extensive base-pairing between nucleotides from the constant and variable regions.
